# Dense Functional and Molecular Readout of a Circuit Hub in Sensory Cortex

**DOI:** 10.1101/2021.02.23.432355

**Authors:** Cameron Condylis, Abed Ghanbari, Nikita Manjrekar, Karina Bistrong, Shenqin Yao, Zizhen Yao, Thuc Nghi Nguyen, Hongkui Zeng, Bosiljka Tasic, Jerry L. Chen

## Abstract

Information processing in the neocortex is carried out by neuronal circuits composed of different cell types. Recent census of the neocortex using single cell transcriptomic profiling has uncovered more than 100 putative cell types which subdivide major classes of excitatory and inhibitory neurons into distinct subclasses. The extent to which this molecular classification predicts distinct functional roles during behavior is unclear. Here, we combined population recordings using two-photon calcium imaging with spatial transcriptomics using multiplexed fluorescent *in situ* hybridization to achieve dense functional and molecular readout of cortical circuits during behavior. We characterized task-related responses across major transcriptomic neuronal subclasses and types in layer 2/3 of primary somatosensory cortex as mice performed a tactile working memory task. We find that as neurons are segregated into increasingly discrete molecular types, their task-related properties continue to differentiate. We identify an excitatory cell type, Baz1a, that is highly driven by tactile stimuli. Baz1a neurons homeostatically maintain stimulus responsiveness during altered sensory experience and show persistent enrichment of subsets of immediately early genes including *Fos*. Measurements of functional and anatomical connectivity reveal that upper layer 2/3 Baz1a neurons preferentially innervate somatostatin-expressing inhibitory neurons. We propose that this connection motif reflects a sensory-driven circuit hub that orchestrates local sensory processing in superficial layers of the neocortex.

## Introduction

Cells of the neocortex can be defined based on their molecular composition, the diversity of which is reflected in their transcriptome. The transcriptional profiles observed across this brain region indicate that cortical populations can be hierarchically subdivided into multiple putative transcriptomic cell classes (e.g., GABAergic, glutamatergic), subclasses (e.g., GABAergic Pvalb) and types (e.g., GABAergic Pvalb Vipr2)^1,2^. Even within a single layer of one cortical area, transcriptional diversity remains high^3^. Evidence suggests that this organization may have developmental origins^4,5^, reflect anatomical specificity^6,7^, or physiological properties^8,9^. The extent to which this diversity relates to information encoding during goal-directed behavior is unclear.

The ability to link molecularly-identified neurons with their function during behavior requires monitoring the activity of cell types *in vivo*. Traditional approaches to label cell types using transgenic lines or post-hoc immunohistochemistry are limited to 1-3 molecular markers^10,11^. This has restricted investigations to classes of excitatory and inhibitory neurons at the broadest hierarchical levels of cell type diversity. Recently-developed techniques for multiplexed spatial transcriptomics dramatically increase the number of genes that can be simultaneously identified in tissue^12–16^. Combinatorial expression patterns of multiple genes can then be used to define finer divisions in the transcriptomic taxonomy corresponding to more specific neuronal subclasses and types. Further, spatial profiling of gene expression in intact tissue readily enables dense multi-modal registration of anatomical and functional measurements across neurons within a single sample^17^. In order to ‘crack’ the neuronal circuits underlying behavior, we developed a platform for <u>C</u>omprehensive <u>R</u>eadout of <u>A</u>ctivity and <u>C</u>ell Type <u>M</u>arkers (CRACK) that combines *in vivo* two photon calcium imaging with post-hoc multiplexed fluorescent *in situ* hybridization. We applied the CRACK platform to study the function of newly identified cell types in layer 2/3 (L2/3) of primary somatosensory cortex (S1).

### CRACK platform

The CRACK platform employs a multi-area two-photon microscope^18^ configured to perform simultaneous population calcium imaging across multiple tissue depths, providing 3D spatial information of neuron location for later post-hoc identification (**Fig. 1a**, **Supplementary Fig. 1, Supplementary Video 1**). Following functional *in vivo* experiments, tissue encompassing the imaged volume is sectioned parallel to the imaging plane. The tissue is embedded in hydrogel and cleared^19^ in order to facilitate labeling of mRNA transcripts using hybridization chain reaction fluorescence *in situ* hybridization (HCR-FISH)^20^ and confocal imaging. Since HCR-FISH is a DNA-based labeling strategy, probes for different mRNA transcripts are labeled, imaged, and then stripped using DNAse across multiple rounds. In order to re-identify and register *in vivo* neurons across multiple rounds of HCR-FISH, we dedicate one imaging channel (561) to repeated labeling and imaging of transcripts of the red genetically encoded calcium indicator, RCaMP1.07, used for functional imaging^21^ (**Supplementary Fig. 2, Supplementary Note S.1**). Other imaging channels are used for labeling cell type-specific markers (**Supplementary Table 1**).

**Figure 1.**
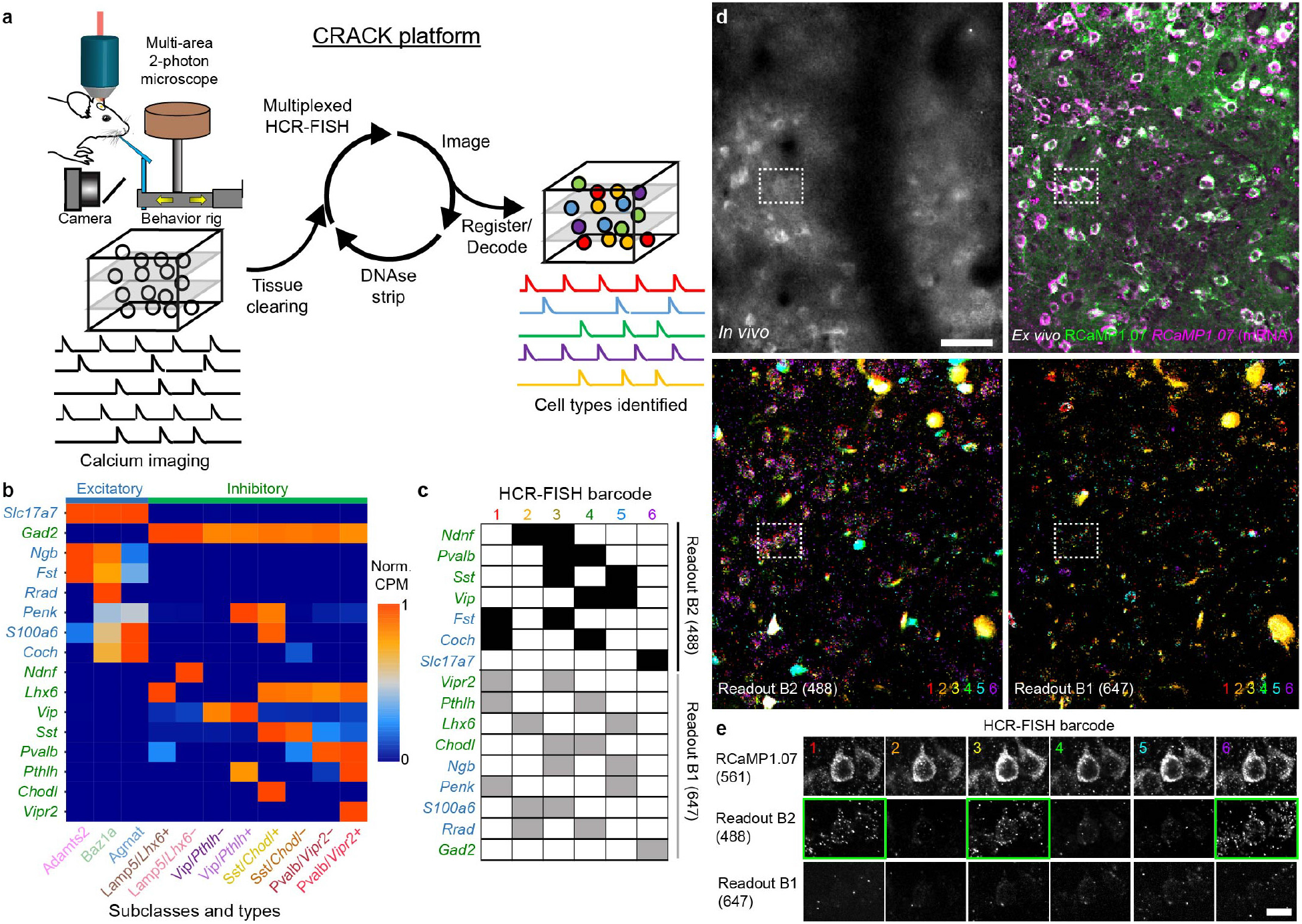
Multiplexed identification of transcriptomic cell subclasses and types in functionally imaged neurons. **a,** Schematic of the CRACK platform. **b,** Expression patterns of genes selected to identify L2/3 S1 excitatory (blue) and inhibitory (green) cell subclasses and types. **c,** Barcode scheme for multiplexed HCR-FISH of selected genes. One set of genes is labeled using B2-488 hairpins while another set of genes is labeled with B1-647 hairpins. **d,** Registration of *in vivo* calcium imaged neurons to *ex vivo* tissue section across multiple rounds of HCR-FISH. Upper left panel shows *in vivo* two-photon images of RCaMP1.07-expressing neurons. Upper right panel shows *ex vivo* confocal images of re-identified neurons showing endogeneous RCaMP1.07 fluorescence following tissue clearing (green) and following HCR-FISH of *RCaMP1.07* transcripts (magenta). Overlay of B2-488 (lower left) and B1-647 (lower right) readout channels across all HCR-FISH barcode rounds. **e,** Decoding of *in vivo* imaged neuron (dotted rectangle in **d**) expressing *Fst* and *Slc17a7*. Individual HCR-FISH rounds are shown with *RCaMP1.07* expression for registration along with B2-488 and B1-647 readout channels. Positive readouts are identified with green rectangle. Scale bars: 50 μm, **d** ; 20 μm, **e**.

While small numbers of genes can be read out through multiple rounds of sequential staining, a barcode readout scheme provides high read depth (100-1000 genes) in an error robust manner. Using barcode readouts to decode arbitrary gene sets relies on single-molecule mRNA resolution which is sensitive to image registration errors and has only been demonstrated in thin tissue sections (<40μm)^12,16^. To obviate the need for single-molecule mRNA-resolution registration so that larger volumes of tissue (150-300μm) can be imaged and analyzed, we programmed our barcode for cellular-resolution readout. This approach relies on prior knowledge of expression patterns to select genes with non-overlapping expression at the cellular level for each imaging channel and hybridization round. Through this, binary decoding at each round occurs at the cellular level rather than the mRNA level. This approach is highly compatible with identifying molecular cell types that are defined by non-overlapping gene expression patterns.

To characterize the hierarchical organization of cell type diversity in L2/3 of S1, we analyzed single cell RNA sequencing (scRNAseq) data from S1 acquired as part of a larger study of the molecular diversity of the isocortex^22^. In this survey, L2/3 pyramidal neurons in S1 were observed to be segregated into three molecular cell types (Adamts2, Baz1a, and Agmat) based on combinatorial expression patterns (**Supplementary Fig. 3**). Excitatory neurons in L2/3 show both cell-type-specific and area-specific gene expression patterns. When comparing S1 cells types to those in primary visual (V1) and anterior lateral motor (ALM) cortex, Baz1a and Agmat cells showed similarity to L2/3 cell types identified in V1 and ALM, whereas Adamts2 cells were present in V1 but not ALM^1^. Molecularly-defined excitatory cell types have been associated with distinct laminar expression patterns or inter-area projection targets^3,4^. We examined whether cell type markers potentially correspond to non-overlapping populations of S1 neurons that project to secondary somatosensory cortex or primary motor cortex^23^. HCR-FISH of retrograde labeled L2/3 neurons indicate that the molecular identities of these transcriptomic cell types do not necessarily correspond to these two projection classes (**Supplementary Fig. 4**).

Inhibitory neuron cell types in S1 were shared with other cortical areas and found to be hierarchically organized^22^. While the major non-overlapping inhibitory subclasses (Lamp5, Pvalb, Sst, Vip) have each been investigated at the broadest level^24,25^, further divisions of these subclasses have not been investigated during task behavior. Thus, we sought to select gene markers that could define the next level of transcriptional subdivision (**Supplementary Fig. 5**). Lamp5 neurons were subdivided into two additional subclasses based on co-expression of either LIM homeobox 6 (*Lhx6*) or neuron derived neurotrophic factor (*Ndnf)*. Pvalb neurons were subdivided based on expression of vasoactive intestinal peptide receptor 2 (*Vipr2*). Sst neurons were subdivided based on expression of chondrolectin (*Chodl*), and Vip neurons were subdivided based on expression of parathryroid hormone–like hormone (*Pthlh*). From this information, we devised a barcode scheme for detection of 16 mRNA species across 6 rounds of staining to resolve 11 cell subclasses or types (3 excitatory, 8 inhibitory) for functional characterization (**Fig. 1b-e**).

### Task encoding across excitatory types

To identify functional differences between transcriptomically-defined cell populations in L2/3 of S1, we performed two-photon calcium imaging on expert wild type mice (*n* = 7) performing a head-fixed whisker-based delayed non-match to sample (DNMS) task^26^ (**Supplementary Fig. 6**). In this context-dependent sensory processing task, a motorized rotor is used to deflect multiple whiskers in either an anterior or posterior direction during an initial ‘sample’ and a later ‘test’ period, separated by a 2s delay (**Fig. 2a**). During the delay period and the inter-trial interval, the rotor was withdrawn to prevent whisker-rotor contact. Behavior was reported as ‘go/no-go,’ in which animals licked on ‘go’ trials for a water reward (‘hit’) when the presented sample and test stimulus were non-matching and withheld licking on ‘no go’ trials (‘correct rejection’) when the presented sample and test stimulus were matching. Misses on go trials were not rewarded, and false alarms on no-go trials were punished with an air puff and a time-out period. In addition to two-photon calcium imaging, high-speed videography was performed to monitor whisking behavior.

**Figure 2.**
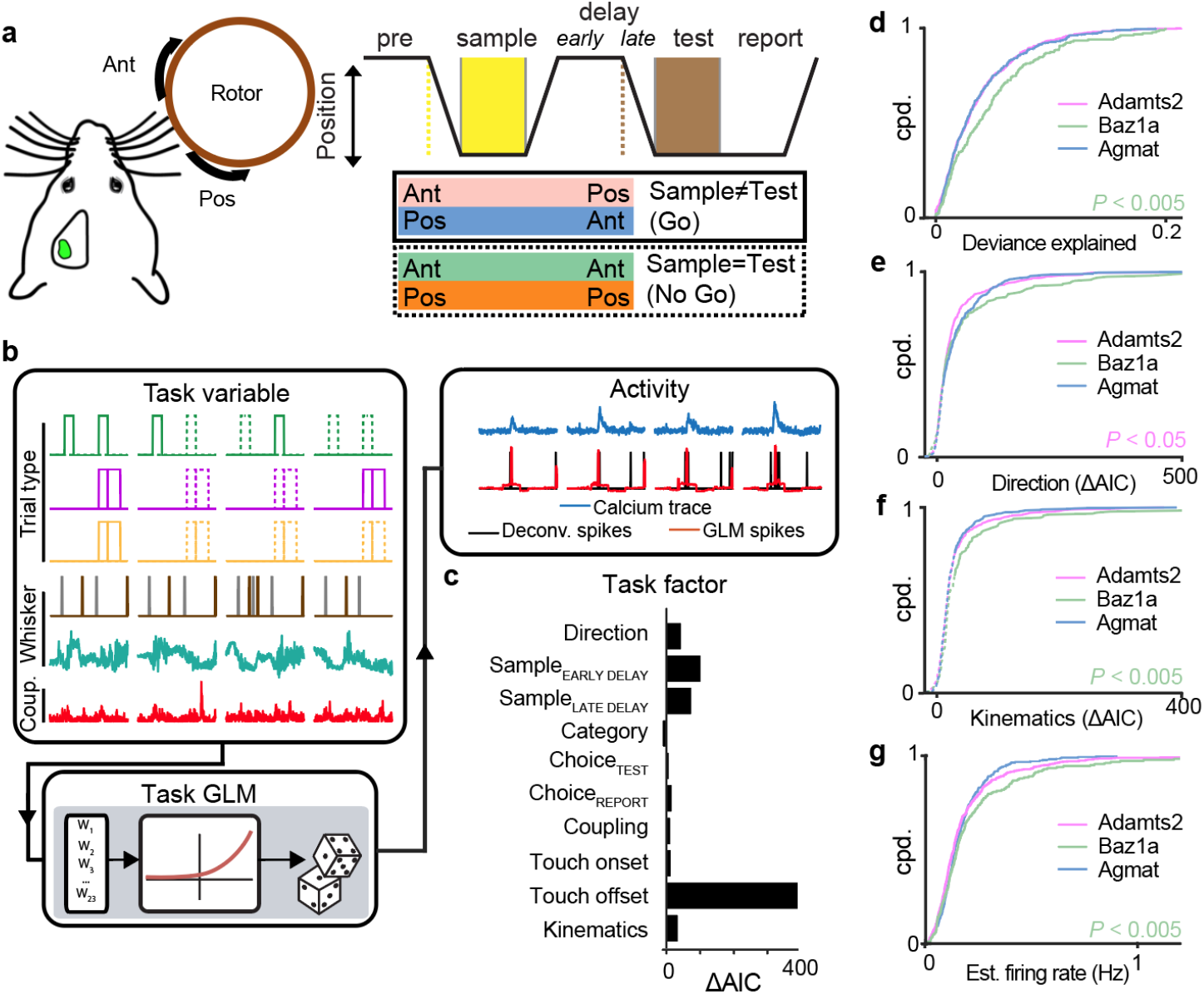
Task encoding across L2/3 excitatory cell types. **a,** Schematic of whisker-based delayed non-match to sample behavioral task. **b,** Encoding of activity in individual neurons using a GLM composed of task variables including trial type information, population activity coupling, and whisker kinematics. **c,** Strength of groups of task variables were determined by comparing full and partial GLM fits (ΔAIC) for individual task factors. Large, positive ΔAIC indicates neuronal encoding of the individual task factor. **d-g**, Cumulative probability distributions of full model deviance explained (**d**), encoding strength of stimulus direction (**e**), encoding strength of whisker kinematics (**f**), and estimated firing rate (**g**) across the three excitatory cell types. (**d-f**, Mann Whitney *U* test; **g**, one-tailed Student’s *t*-test). In **e-f**, solid and dotted lines correspond to significant (*P* < 0.01) and non-significant encoding strengths via χ^2^ test. *n* = 1107 neurons from 7 animals.

We previously reported diverse task-related responses in L2/3 of S1 during the DNMS task^26^. To characterize task-related responses for each recorded cell in a more comprehensive manner, we fit a generalized linear model (GLM) to each neuron’s estimated spiking activity against a range of ‘task variables’^27^ (**Fig. 2b**, **Supplementary Fig. 8-9, Supplementary Note S.2**). Task variables representing a related feature were grouped into ‘task factors’(i.e. stimulus direction, trial category, etc.). The ability for a neuron to encode a particular task factor was determined by calculating the difference in the Akaike Information Criterion (ΔAIC) between a full model and a partial model excluding task variables representing that task factor. A positive ΔAIC value indicates reduced fit quality from the full to the partial model, revealing that the excluded task factor in the partial model is an important contributor to the modeled neuron’s activity. Thus, we interpret significant, positive ΔAIC values to indicate neuronal encoding of the excluded task factor (**Fig. 2c, Supplementary Fig. 10, Supplementary Note S.3**). We analyzed 10 task-related factors. Six of the ten task factors were defined by trial type information. This included information related to the direction of the task stimulus (direction), trial category defined by the combination of the sample and test stimulus (category), and the animal’s choice during the test (choice_TEST_) and report period (choice_REPORT_). While our previous study found no evidence of sustained activity in S1 during the delay period, we included task factors representing the sample stimulus at later points in the trial (sample encoded early in the delay period, sample_EARLY DELAY_; Sample information late in the delay period, sample_LATE DELAY_). Another set of task factors describing whisker movement and tactile-object interactions were derived from video analysis of whisker tracking and included whisker-object touch onset (touch onset), whisker-object touch offset (touch offset), and whisker kinematics (kinematics). A final task factor was derived from the activity of all other simultaneously recorded neurons to assess the level of coupling the neuron had with overall network activity (coupling)^28^.

We first compared differences in task encoding across the three excitatory cell types. We observed that Baz1a showed the best overall GLM fit (**Fig. 2d**). In particular, Baz1a neurons more strongly encoded whisker kinematics compared to the other two excitatory cell types (*P* < 0.005, Mann Whitney *U* test; **Fig. 2f**). In contrast, Adamts2 neurons more weakly encoded stimulus direction and touch offset (direction: *P* < 0.05; touch offset: *P* < 0.02, Mann Whitney *U* test; **Fig. 2e, Supplementary Fig. 11**,) while Agmat neurons more strongly encoded choice_REPORT_ (*P* < 0.05, Mann Whitney *U* test). Baz1a neurons also show overall higher firing rates (*P* < 0.005, one-tailed Student’s *t-*test) and response reliability to sample and test stimuli (**Fig. 2g, Supplementary Fig. 12**). Taken together, these findings suggest that Baz1a neurons represent a highly active, sensory-driven L2/3 excitatory cell type.

### Persistent stimulus activity and *Fos* expression in Baz1a neurons

Highly active, sensory-driven L2/3 S1 neurons have been previously observed to exhibit high expression of the immediate early gene, *Fos*^29,30^. Single-cell RNAseq analysis in naïve, untrained mice shows that while all cell types express some number of immediate early genes (IEGs), Baz1a neurons show consistent enrichment of *Fos* along with other subsets of IEGs (**Fig. 3a**). Previous studies observed *Fos* expression to be dynamic and driven by experience-dependent plasticity^31^. However, due its conserved and elevated expression in Baz1a cells across naïve animals, we speculated that *Fos* and other IEGs may be stably expressed in Baz1a neurons. To confirm this and address how it relates to neuronal function, we extended the CRACK platform to track *Fos* expression and stimulus activity during altered sensory experience using transgenic fosGFP mice^32^along with virally co-expressed Rcamp1.07 in S1 (*n* = 3; **Fig. 3b,c**). *Ex vivo* HCR-FISH confirmed that high fosGFP fluorescence corresponded with higher *Fos* mRNA and that *GFP* mRNA was also enriched in Baz1a neurons (**Fig. 3d, Supplementary Fig. 13**).

**Figure 3.**
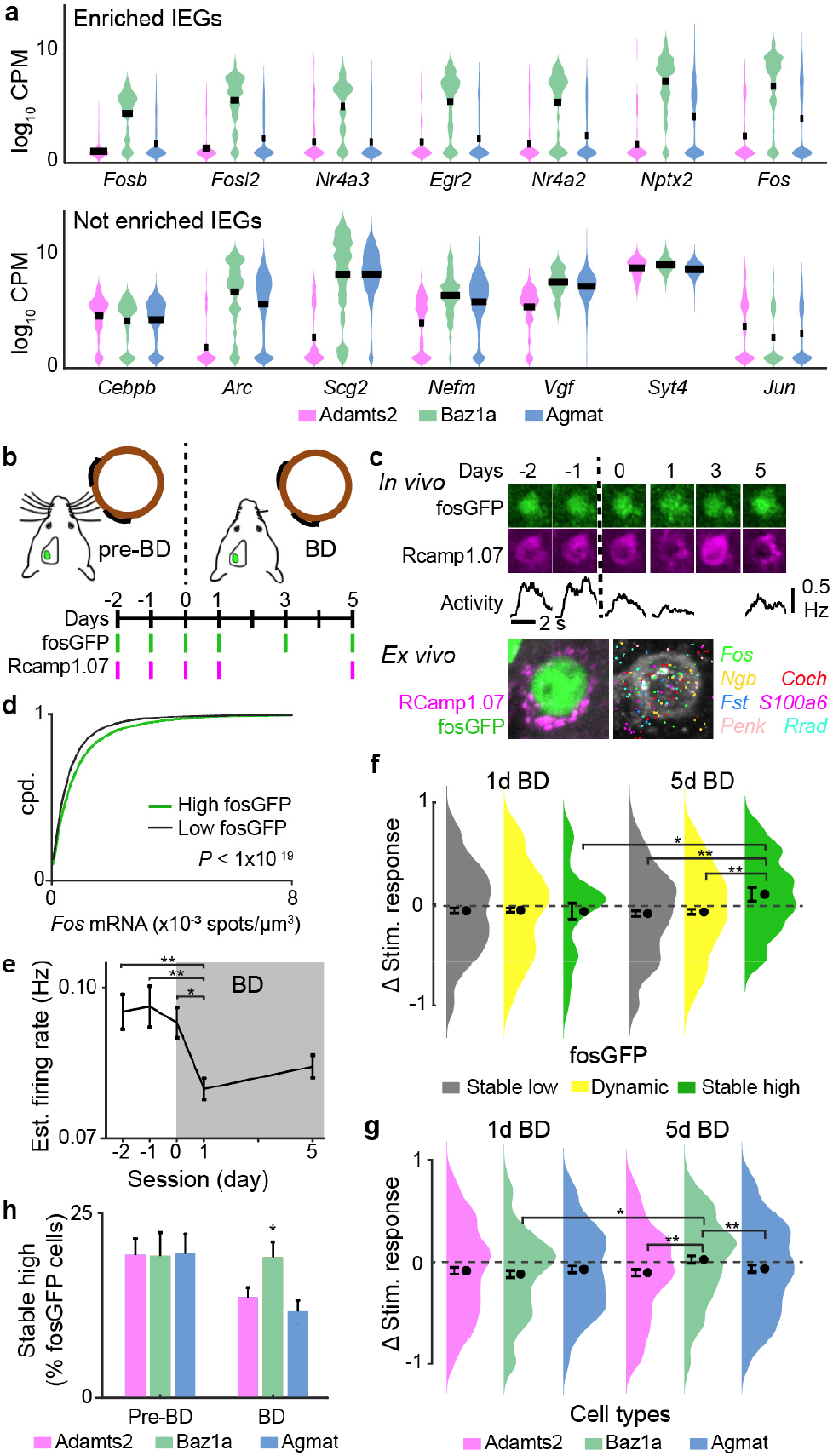
Persistent IEG expression and homeostatic plasticity in Baz1a neurons. **a,** Example of immediate early genes selectively enriched (top) and not enriched (bottom) in Baz1a cells. **b,** Time course of bilateral whisker deprivation (BD) experiment. **c,** Example of Baz1a neuron with stable high fosGFP expression across *in vivo* imaging sessions (top). Average stimulus responses during calcium imaging are shown (middle). Post-hoc identification of neuron and HCR-FISH for select genes are shown (bottom). **d,** HCR-FISH *Fos* spot density in high (1.2-fold above background) and low fluorescent fosGFP cells (two-tailed Student’s *t*-test). **e,** Mean stimulus-evoked activity before and after BD across functionally imaged neurons (one-way ANOVA with post-hoc multiple comparison test, *n* = 2569 cells from 3 animals). **f,** Change in stimulus-evoked responses before BD versus at 1 day or 5 days BD across neurons with stable low, dynamic, and stable high fosGFP expression (two-tailed Student’s *t*-test, *n* = 790 cells from 3 animals). **g,** Change in stimulus-evoked responses before BD versus at 1 day or 5 days BD across excitatory cell types (χ^2^ test, *n* = 511 cells from 3 animals). **h,** Fraction of fosGFP neurons with stable high expression across all pre-BD sessions (day −2, −1, 0) and across all BD sessions (1, 3, 5) for excitatory cell types (two-tailed Student’s *t*-test, *n* = 3753 cells from 3 animals). (* *P* < 0.05, ** *P* < 0.005). Error bars = s.e.m, **e-g**; s.d. from bootstrap analysis, **g**.

FosGFP and sensory-evoked calcium responses were tracked before and after 5 days of bilateral whisker deprivation (BD). During BD, the principal whisker corresponding to the imaged S1 barrel column was trimmed to a minimum length so that stimulus-evoked activity could still be tracked. Overall, BD resulted in a decrease in stimulus-evoked activity after 1 day followed by a slow homeostatic compensation after 5 days similar to as previously observed^33^ (*P* < 0.0002, one-way ANOVA, post-hoc multiple comparison test; **Fig. 3e**). We first asked how fosGFP expression related to functional changes during BD. Cells were divided into 3 groups based on fosGFP expression: 1) stable low fosGFP expression across all imaging sessions; 2) stable high fosGFP expression across all imaging sessions, and; 3) dynamic fosGFP expression between at least one imaging session (**Fig. 3f**). All groups showed decreased stimulus-evoked activity after 1 day of BD. However after 5 days, responses in stable low and dynamic fosGFP neurons remained depressed while stable high fosGFP neurons exhibited an enhancement in sensory response magnitude compared to pre-BD conditions (*P* < 0.05, Student’s *t*-test).

*Ex vivo* cell type identification revealed that similar fractions of stable high fosGFP neurons were observed across all excitatory cell types before deprivation. However, during BD, there was increased fosGFP turnover in Adamts2 and Agmat neurons while the fraction of stable high fosGFP cells remained unchanged in Baz1a neurons (*P* < 0.05, χ^2^ test; **Fig. 3h**). Functionally, all three cell types showed reduced stimulus activity after 1 day BD while only Baz1a neurons showed recovery after 5 days of BD (*P* < 0.005, Student’s *t*-test; **Fig. 3g**). Taken together, this demonstrates that Baz1a neurons are molecularly and functionally distinct in exhibiting stable enrichment of select IEGs and maintaining stimulus responsiveness during altered sensory experience.

### Task encoding in inhibitory classes and subdivisions

We next compared task encoding in three of the major classes of inhibitory neurons (Pvalb, Sst, Vip). Lamp5 neurons were excluded from analysis due to their low numbers captured in the data set (**Supplementary Table 2**). Overall, Pvalb neurons exhibited the weakest coding of tactile-related features such as direction, touch onset, kinematics, as well as sample_LATE DELAY_ (**Fig 4a-c, Supplementary Fig. 14**). However, the high firing rates of Pvalb neurons and associated difficulties in reliably inferring spiking activity in this subclass by calcium imaging may underestimate the strength of GLM-derived task responses^11,34^ (**Supplementary Note S.2**). We therefore focused our analysis on Sst and Vip neurons.

**Figure 4.**
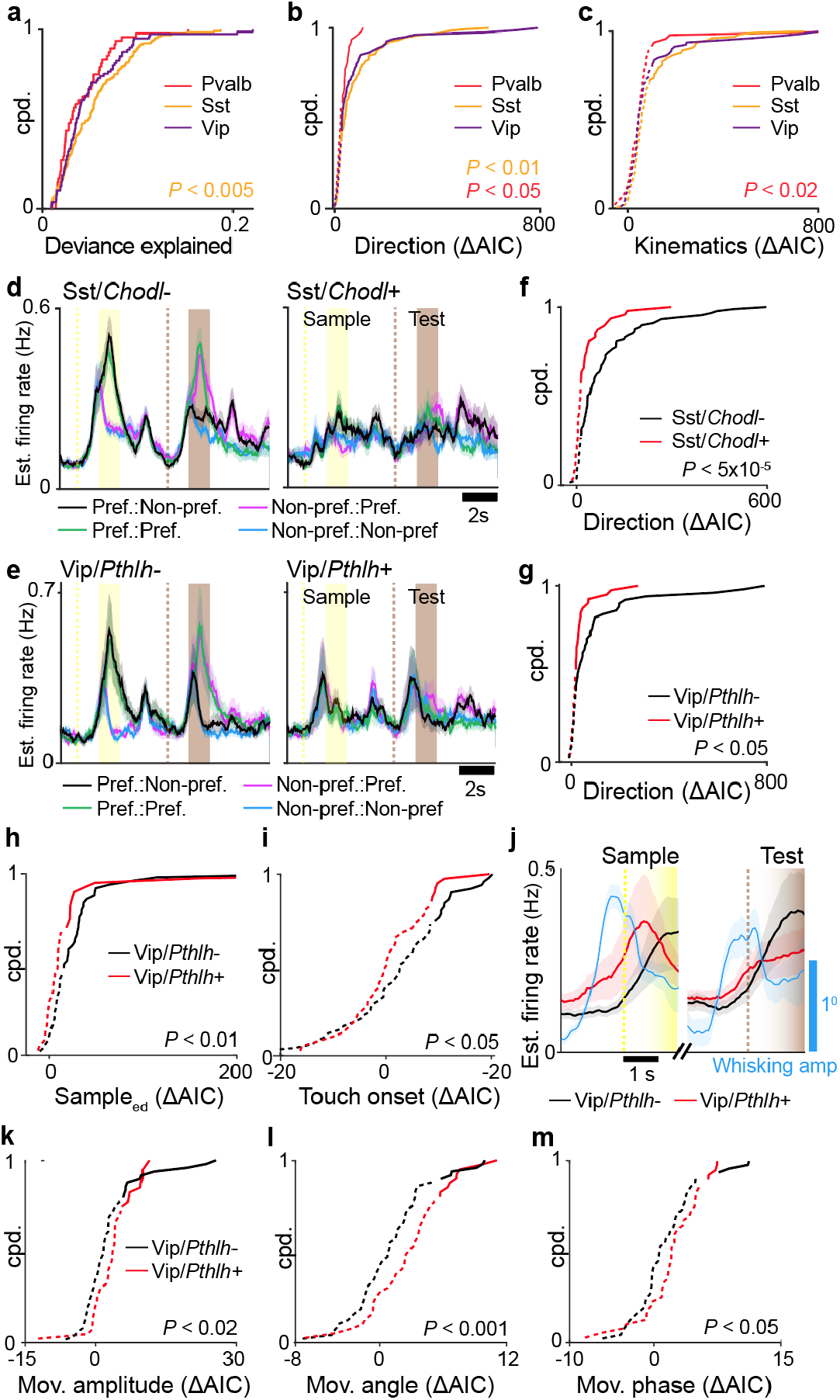
Task encoding across L/3 inhibitory subclasses. **a-c,** Cumulative probability distributions for full model deviance explained (**a**), encoding strength of stimulus direction (**b**), and encoding strength of whisker kinematics (**c**) for three major inhibitory cell types (Mann Whitney *U* Test). **d, e,** Estimated firing rate responses to preferred stimulus direction for Sst subclasses (**d**) and Vip subclasses (**e**). **f, g,** Cumulative probability distribution of ΔAIC for task factor encoding direction for Sst subclasses (**f**) and Vip subclasses (**g**) (Mann Whitney *U* Test). **h, i,** Cumulative probability distribution of ΔAIC for tasks factors encoding sample_EARLY DELAY_ (**h**) and touch onset (**i**) for Vip subclasses (Mann Whitney *U* Test). **j,** Estimated firing rate for Vip subclasses along with mean whisking amplitude aligned to whisker-rotor touch onset preceding sample and test periods. **k-m.** Cumulative probability distribution of ΔAIC for task factors encoding free whisking amplitude (**k**), angle (**l**) and phase (**m**) for Vip subclasses (Mann Whitney *U* Test). In **b-c, f-g, m-f**, solid and dotted lines correspond to significant (*P* < 0.01) and non-significant encoding strengths via χ^2^ test. Shaded region in **a**, **c** corresponds to s.e.m. *n* = 48 Pvalb cells, 47 Sst/*Chodl*+ cells, 88 Sst/*Chodl*-cells, 40 Vip/*Pthlh*+ cells, 49 Vip/*Pthlh*-cells from 7 animals.

We sought to investigate if more task-related differences emerge as inhibitory classes are further subdivided into more discrete transcriptomic subclasses or types. As a major class, Sst showed the best overall fit to the GLM (*P* < 0.005, Mann-Whitney *U* test; **Fig. 4a**) and strongly encoded stimulus direction (*P* < 0.05, Mann-Whitney *U* test; **Fig. 4b**). We asked whether stimulus direction was encoded similarly in subclasses of Sst neurons. Sst/*Chodl*+ neurons express nitric oxide synthase (*Nos1*)^35^, display long-range axonal projection patterns^36,37^, and are active during slow wave sleep^38^. We find that during the DNMS task, Sst/*Chodl*+ encoded direction more weakly compared to Sst/*Chodl*- neurons (*P* < 5 × 10^−5^, Mann Whitney *U* test; **Fig. 4d,f, Supplementary Fig. 15a**). These results demonstrate that stimulus-driven responses are specific to subsets of Sst neurons that do not express *Chodl.*

We next compared task differences between Vip/*Pthlh*+ and Vip/*Pthlh*- neurons (**Fig. 4e**). Vip neurons belonging to the *Pthlh*+ subclass co-express choline acetyltransferase (*Chat*) and calretinin, also known as calbindin-2 (*Calb2*), typically have bipolar morphologies, and preferentially target Sst neurons^37,39–41^. Vip neurons belonging to the *Pthlh*- subclass co-express synuclein gamma (*Sncg*) and cholecystokinin (*Cck*), have multipolar and basket cell morphologies, and preferentially target Pvalb neurons^37,42^. We found that Vip/*Pthlh*- neurons more strongly encoded direction, sample_EARLY DELAY_, and onset than Vip/*Pthlh*+ neurons (direction: *P* < 0.05; sample_EARLY DELAY_: *P* < 0.01; onset: *P* < 0.05, Mann-Whitney *U* test; **Fig. 4g-i**, **Supplementary Fig. 15b**). Analysis of neuronal firing with respect to touch onset at the beginning of the sample and test period neurons showed elevated firing for Vip/*Pthlh*+ neurons preceding touch onset, which correlated with an anticipatory increase in whisking amplitude (**Fig. 4j**). This pre-touch activity suggested that Vip/*Pthlh*+ neurons are driven by free whisking behavior. To disentangle movement-related from tactile-related whisker responses, we fit neuronal activity to a GLM with whisker kinematic variables using only time periods prior to touch onset during the pre-stimulus and delay period. We found that Vip/*Pthlh*+ neurons more strongly encoded whisker amplitude, angle, and phase task factors during free whisking periods compared to Vip/*Pthlh*- neurons (amplitude: *P* < 0.02; angle; *P* < 0.001; phase: *P* < 0.05, Mann-Whitney *U* test; **Fig. 4k-m**). These results indicate that the activity related to whisking behavior previously identified in Vip neurons^41^ is specific to Vip/*Pthlh*+ neurons. Overall, these findings demonstrate further segregation of functional task properties in inhibitory neurons with increasing transcriptomic subdivisions.

### Network interactions between major subclasses and types

The ability to simultaneously record across all identified cell subclasses and types enables a comprehensive characterization of cell type-specific network structures that underlie coding of task information. Non-negative matrix factorization across varying ranks can be used to capture population dynamics across distinct functional subpopulations (**Supplementary Note S.4**). Neurons that exhibit strong population coupling with increasing ranks suggest functional relationships with multiple subpopulations. Compared to other excitatory neurons, Baz1a neurons consistently showed higher coupling across increasing coupling ranks indicating that they are highly integrated into the local L2/3 network (*P* < 0.02, *F_2,6,_* repeated measures ANOVA; **Fig. 5a**). To investigate how this coupling relates to specific cell type populations, we constructed a GLM that included all previously described task variables while subdividing the activity of simultaneously recorded neuronal activity into different “coupling factors” according to Adamts2, Baz1a, Agmat, Pvalb, Sst, and Vip transcriptomic populations (**Fig. 5b**). For a modeled neuron, the ΔAIC for each cell type coupling factor constituted a measure of “functional connectivity” between that neuron and other simultaneously recorded cell types that could reflect either positive or negative noise correlations (**Supplementary Fig. 16**). From these measures, a directional weighted network graph can be constructed composed of the six cell populations as nodes and functional connectivity as edges. Task-specific networks were generated by selecting for neurons with significant ΔAIC (*P* < 0.01, χ^2^ test) for a given task factor (**Fig. 5c**). This allowed us to assess the degree of interactions between cell populations encoding different task factors.

**Figure 5.**
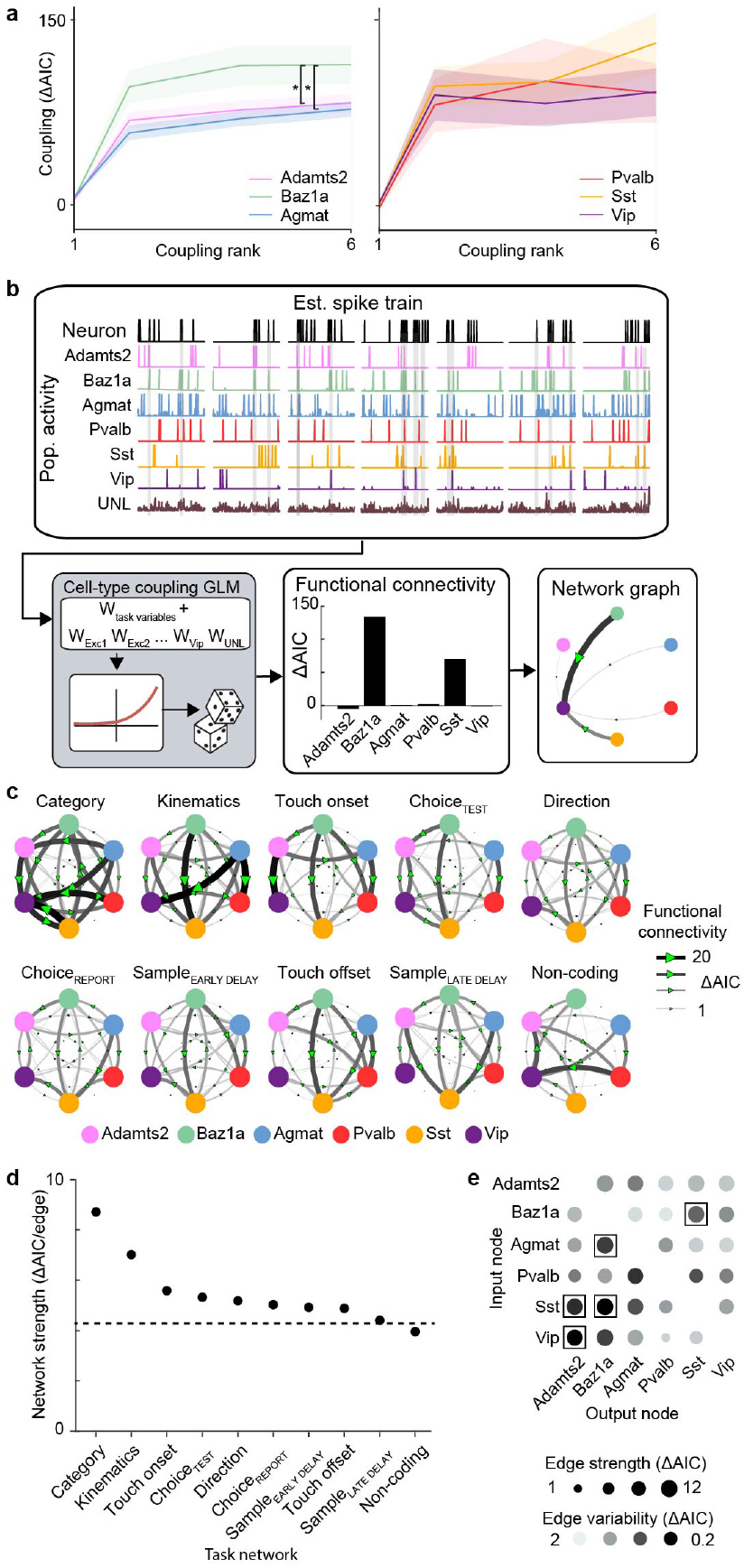
Cell type functional connectivity across task networks. **a,** Strength of coupling factor encoding across varying coupling ranks (\**P* < 0.02, repeated measures ANOVA test, *F2,6*). **b,** Schematic of network analysis for example neuron. Activity from simultaneously recorded excitatory types and inhibitory subclasses were used to construct a cell type coupling GLM. Cell type coupling factors were calculated as measures of functional connectivity to build network graphs of interactions between cell types. **c,** Task-specific networks generated by selecting for neurons with significant encoding for a given task factor in the task GLM. Networks are sorted according to average edge strength. **d,** Network strength across task networks. Dotted line corresponds to strength of shuffled network. **e,** Strength and variability of functional connectivity in network edges across task networks. Network edges with significantly high strength and low variability are indicated with a box (*P* < 0.05, permutation test). Shaded region in **a** corresponds to s.e.m. *n =* 1996 neurons, Direction; 1374 neurons, sample_EARLY DELAY_; 1076 neurons sample_LATE DELAY_; 360 neurons, Category; 623 neurons, choice_TEST_; 830 neurons, choice_REPORT_; 898 neurons, touch onset 1033 neurons, touch offset; 864 neurons, kinematics; 273 neurons, non-coding from 7 animals.

We observed different network patterns across task factors. All task factor networks exhibited cell type-specific functional connectivity weights that were greater than chance (with the exception of the network containing non-coding neurons, which exhibited random connectivity) (*P* < 0.05, bootstrap test; **Fig 5d**). Functional connectivity was strongest amongst neurons encoding category and whisker kinematics, suggesting that encoding of these task variables involves a high degree of local interactions (**Supplementary Fig. 17a)**.

We further investigated the structure of these networks. For each cell type node, we used the input edge strengths to determine how other cell populations influence the activity of the measured node and the output edge strength to measure how the measured node influences the activity of other cell populations (**Supplementary Fig. 17b**). Overall, we observed that inhibitory neurons were more likely to be influenced by network activity patterns than excitatory neurons. Sst neurons exhibited the highest input node strength across all task conditions (*P* < 0.05, bootstrap test). This is in line with evidence suggesting that Sst neurons follow local network activity^25^. In contrast, excitatory neurons had a greater influence on other cell types with Baz1a cells showing high output node strength in 7 out of the 9 task factor networks (*P* < 0.05, bootstrap test). This suggests that Baz1a cells orchestrate a majority of local L2/3 activity patterns.

Given the differences in node strengths across task factor networks, we asked whether functional connectivity between any two cell types reflected a dynamic relationship that depended on the task factor network or represented a stable motif that is potentially intrinsic to the underlying circuitry. To address this, we measured the overall strength of each connection by calculating the mean edge weight across task factor networks. The stability of this connection was reported as the coefficient of variation of the edge weight across task factor networks (**Fig. 5e**). The majority of connections exhibited variability between task factor networks that were equivalent to chance levels, suggesting that the strength of those connections are dynamic and depend on the encoded task factor. However, a subset of connections (Adamts2→Vip, Adamts2→Sst, Agmat→Baz1a, Baz1a↔Sst; output node→input node) were consistently strong and stable across task factor networks, suggesting that they represent intrinsic functional motifs between specific cell populations (*P* < 0.05, permutation test). These findings suggest that processing of task information occurs within local circuits that are composed of both dynamic and fixed interactions between transcriptomic cell populations.

### Cell-type specific tracing confirms intrinsic functional connectivity

The observed functional connections that persisted across task networks could be explained by cell type-specific synaptic connections. Trans-monosynaptic rabies tracing enables input mapping to specific cell types but requires genetic access for conditional infection. Since transgenic lines for the three excitatory cell types are not available, we focused on input patterns to Sst and Vip inhibitory classes. Since Baz1a neurons showed stable functional connectivity with Sst neurons but not Vip neurons, we sought to compare Baz1a synaptic connectivity between these two inhibitory classes. Using Sst-IRES-Cre (*n* = 4) and Vip-IRES-Cre (*n* = 4) mice^43^, L2/3 Sst and Vip starter cells were first labeled using a Cre-dependent AAV expressing TVA, CVS-N2cG, and dTomato, followed by delivery of the EnvA-pseudotyped CVS-N2c(ΔG) rabies virus expressing histone-GFP^44^ (**Fig. 6a,b**). We first examined the sublaminar distribution of histone-GFP positive inputs (nGFP+) to Sst and Vip cells across L2/3. Overall, Sst and Vip neurons received greater number of inputs from cells located in deeper L2/3 (>200μm below the pia) compared to superficial L2/3. However, we found Sst neurons received more of their inputs from superficial L2/3 neurons compared to Vip neurons (Sst: 28.9±1.6%; Vip: 21.5±2.8%; mean±s.e.m.; *P* < 0.02, one-tailed Student’s *t*-test; **Fig. 6c**). Multiplexed HCR-FISH from the CRACK platform was performed to identify cell type of nGFP+ input neurons. The overall density Baz1a inputs was greater for Sst neurons compared to Vip neurons (Sst: 12.2±1.3%; Vip: 8.1±1.5%; *P* < 0.01; one-tailed Student’s *t*-test; **Fig. 6d**). This difference was greatest among cells in superficial L2/3 (Sst: 21.1±1.7%; Vip: 16.9±1.6%; *P* < 0.05; one-tailed Student’s *t*-test; **Fig. 6e**). These results reveal differential sublaminar organization of the input patterns to Sst and Vip neurons and identify a cell type specific synaptic connection between superficial L2/3 Baz1a and Sst neurons that is consistent with functional connectivity measurements.

**Figure 6.**
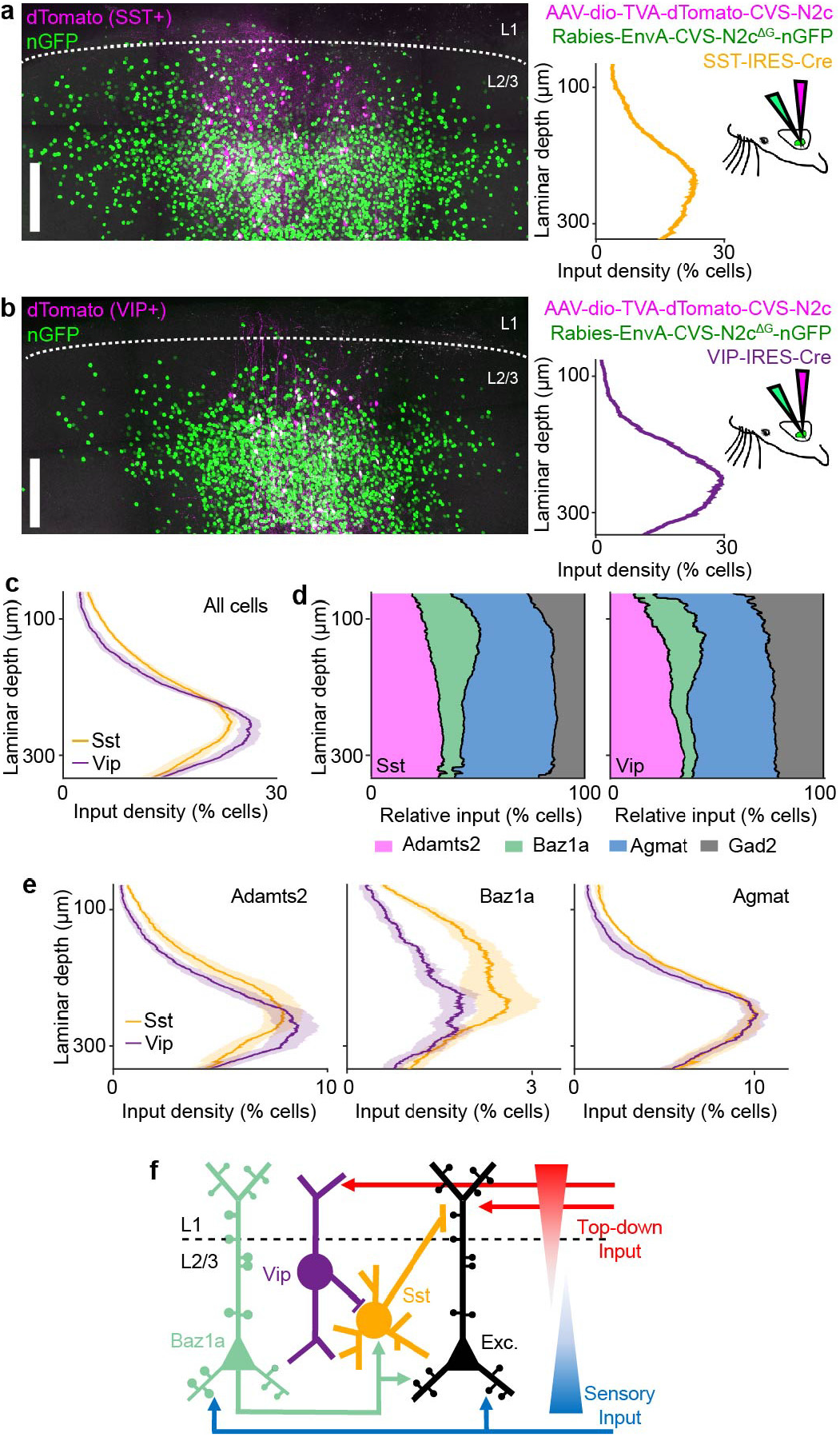
Upper layer Ba1za neurons target Sst neurons. **a-b,** Example of cell type-specific trans-monosynaptic tracing in Sst-IRES-Cre (**a**) and Vip-IRES-Cre (**b**) mice. Left panels show confocal images from coronal sections with starter cells (magenta) and nGFP+ input neurons (green). Right panels show sublaminar distribution of input density as a function of depth from left images along with injection scheme. **c**, Average sublaminar somatic density distribution of inputs across L2/3 for Sst and VIP neurons. **d**, Relative proportion of excitatory cell types and *Gad2*+ inhibitory neurons as a function of laminar depth for Sst and Vip input neurons. **e,** Sublaminar density distribution for excitatory cell types as a function of laminar depth for Sst and Vip input neurons. **f,** Circuit model of L2/3 illustrating cell-type-specific connectivity between Vip, Sst, Baz1a, and other local excitatory neurons. Sensory-driven activation of Baz1a neurons can inhibit top-down inputs via activation of Sst neurons, biasing integration of sensory and recurrent inputs in local excitatory neurons. Shaded region in **c,e** corresponds to s.e.m. *n =* 16 slices, 33,957 neurons from 4 Sst-IRES-Cre animals, 14 slices; 35,926 neurons 4 Vip-IRES-Cre animals, 14 slices. Scale bars: 100 μm.

## Discussion

In conclusion, we have developed a platform to densely survey the functional and molecular properties of neuronal populations *in vivo* and have applied it to study cell types in L2/3 of S1. Through this, we have shed light on the role of Baz1a neurons in neocortical function. Enriched *Fos* expression suggests that Baz1a neurons are members of a previously described, highly interconnected FosGFP population that operates as a network hub in S1^29^. S1 is important for both tactile feature discrimination as well as sensorimotor integration for object localization^45^. Superficial L2/3 pyramidal neurons are laminarly situated to integrate both top-down motor and associative signals arriving in L1 onto apical dendrites with bottom-up sensory signal arriving from L4 onto basal dendrites^46,47^ (**Fig. 6f**). Basal dendrites also contain highly recurrent, lateral connections between neighboring excitatory neurons^48^. L2/3 Sst neurons predominantly target and inhibit apical dendrite activity^49^. In turn, Vip neurons inhibit Sst neurons to form a disinhibitory circuit motif that enables integration of top-down signals and mediates associative memory formation in L2/3 pyramidal neurons^41,50^. We propose that excitatory connections from superficial Baz1a neurons onto Sst neurons serves to counteract this disinhibitory circuit by driving Sst neurons to inhibit top-down inputs in neighboring pyramidal neurons. This would bias synaptic integration in local L2/3 pyramidal neurons towards bottom-up and recurrent inputs. Showing high responsiveness to sensory stimuli, Baz1a neurons are well poised to respond to sensory stimulation and recruit local circuits for sensory processing. Therefore, these circuit motifs operate complementary to one another, allowing S1 to shift between gating long-range feedback inputs and engaging feedforward computations.

Baz1a neurons are also functionally distinct in their ability to adapt to altered sensory experience by homeostatically maintaining their response to tactile stimuli. Sensory deprivation has been shown to transiently induce changes in IEG expression resulting in experience-dependent plasticity^51^. We speculate that stable expression of *Fos* and other select IEGs in Baz1a cells may prime this cell type to respond to changes in experience through molecular mechanisms that could adjust excitatory-inhibitory balance, synaptic scaling, or intrinsic excitability. This plasticity suggests that Baz1a neurons both act to preserve existing sensory representations in the face of novel experiences and to recruit local circuits for ongoing sensory processing. The presence of cell types in V1 and ALM with similar expression profiles as Baz1a neurons suggest that homologous circuits may exist across neocortical areas. Overall, the findings that we have produced from the CRACK platform provide new insight into how neural computations can be embedded within the molecular logic of neocortical circuits.

## Acknowledgements

We thank O. Gonen, S. Kenyon, G. Shechter, N. Weston, C. Xin for software development, A. Ahrens, G. House, K. Marmon for assistance in image analysis, N. Josephs for advice in statistical analysis, M. Economo, D. Lee, B. Scott, C. Habjan for comments on the manuscript. This work was supported by grants from a NARSAD Young Investigator Grant from the Brain & Behavior Research Foundation, the Richard and Susan Smith Family Foundation, Elizabeth and Stuart Pratt Career Development Award, the Whitehall Foundation, Harvard NeuroDiscovery Center, National Institutes of Health BRAIN Initiative Award (R01NS109965 to J.C., and U19MH114830 to H.Z.), National Institutes of Health New Innovator Award (DP2NS111134), and National Institutes of Health Ruth L. Kirschstein Predoctoral Individual National Research Service Award (F31NS111896) to C.C,.

## Authors’ Contributions

C.C. and J.L.C. designed the study. C.C. performed two-photon imaging and CRACK platform experiments. C.C. and K.B. performed retrograde tracing experiments. S.Y. and C.C. performed rabies tracing experiments. C.C., A.G., N.M., K.B., and J.L.C. performed data analysis. T.N., Z.Y., and B.T. generated and analysed the single cell transcriptomic data supervised by H.Z. C.C. and J.L.C. wrote the paper.

## Declaration of Interests

The authors declare no competing interests.

## Materials & Correspondence

Correspondence to Jerry L. Chen

## METHODS

### Mice

Experiments in this study were approved by the Institutional Animal Care and Use Committee at Boston University, approved by the Allen Institute Animal Care and Use Committee, and conform to NIH guidelines. Behavior experiments were performed using C57BL/6J mice (The Jackson Laboratory). Input mapping experiments were performed using SST-IRES-Cre and VIP-IRES-Cre mice^43^. Binocular deprivation experiments were performed using B6.CgTg(Fos/EGFP)1-3Brth/J mice^32^. All animals were 6-8 weeks of age at time of surgery. Mice used for behavior were housed individually in reverse 12 hour light cycle conditions. All handling and behavior occurred under simulated night time conditions.

### Single cell RNA sequencing analysis

Single cell RNA sequencing data in this study were previously acquired^22^. S1 excitatory and inhibitory subclasses and cell types obtained were identified using an iterative clustering R package hicat (https://github.com/AllenInstitute/hicat) as previously described^1^. To understand the cell type and area differences of L2/3 neurons among S1, V1, and ALM, differentially expressed genes (DEG) identified between all pairs of cell types, all pairs of areas, and all pairs of cell type and area groups (fold change > 4, FDR 0.01, and expressed in >= 40% cells in one group). The average counts per million reads mapped (CPM) values of each DEG within each group were scaled across all the groups to between 0 and 1. A DEG was categorized as cell type specific if its normalized values were greater than 0.5 in at least two areas (or one region if the cluster is specific to one area) within a given cell type, and area-specific if its scaled values were greater than 0.5 in at least two cell types within a given area. DEGs specific to both cell type and area were defined as genes with scaled value greater than 0.5 in only one area and cell type. Immediate early genes enriched in Baz1a neurons were identified as those with statistically higher CPM values compared to Adamts2 and Agmat neurons.

### Animal imaging preparation

For behavior and bilateral whisker deprivation experiments, the genetically encoded calcium indicator RCaMP1.07^21^ was expressed by stereotaxic injection of AAV2/PHP.eb-RCaMP1.07 into S1 (600 nL, ~1 × 10^9^ vg/ml per virus). L2/3 was targeted at 1.1 mm posterior to bregma, 3.3 mm lateral, 300 μm below the pial surface. A 4mm cranial window was implanted over S1 to obtain optical access^52^. A metal headpost was implanted on the skull adjacent to the window to allow for head fixation. One week after window implantation and injections, animals were handled and acclimated to head fixation. In order to identify regions of S1 that correspond to specific whiskers, functional mapping was performed using intrinsic signal imaging.

### Behavioral task

Animals were trained on a head-fixed, whisker-based delayed non-match to sample using equipment and training protocol previously described^26^. Whiskers were trimmed to a single row corresponding to the selected imaging region for high-speed videography. Animals were water deprived throughout training and imaging and only received water by performing the task. Their weight was monitored daily to ensure body weight did not drop below 80% of initial weight. In the task, a motorized rotor was used to deflect whiskers in either an anterior or posterior direction. During the delay period and inter-trial interval, the rotor was withdrawn to prevent whisker-rotor contact. Behavior was reported as ‘go/no-go’ in which animals licked on ‘go’ trials for a water reward (‘hit’) when the presented sample and test stimulus were non-matching and withheld licking on ‘no-go’ trials (‘correct rejection’) when the presented sample and test stimulus were matching. Misses on go trials were not rewarded, and false alarms on no-go trials were punished with an air puff and a time-out period.

### Bilateral whisker deprivation

Awake, head-fixed animals were subjected to passive whisker deflection. Each trial had the following structure: baseline (3s), rotor approach, anterior rotation and whisker deflection (1s), posterior rotation and whisker deflection (1s). For baseline activity and fosGFP measurements, neurons were imaged for two days prior to deprivation (Day −2 and −1). Whiskers were trimmed bilaterally and imaged immediately afterwards (Day 0). The principle whisker was trimmed to a stub ~1-2 mm in length, so that it would not provide sensory information to the animal but could still be deflected by the rotor. Whiskers were re-trimmed every other day. During deprivation, activity measurements were acquired at Day 1 and 5 while fosGFP measurements were acquired at Day 1, 3, and 5.

### Two-photon imaging

Two-photon calcium imaging in expert animals was performed with a custom-built resonant-scanning multi-area two-photon microscope with 16x/0.8NA water immersion objective (Nikon) using custom-written Scope software as previously described^53^. A 31.25 MHz 1040 nm fiber laser (Spark Lasers) was used to excite the red calcium indicator RCaMP1.07. The multi-area two-photon microscope was configured to perform simultaneous imaging at 32.6 Hz frame rate of two imaging planes were separated by ~50 μm in depth. Average power of each beam at the sample was 70-90mW. For each animal, imaging was performed across 8-10 behavioral sessions. For bilateral whisker deprivation experiments, calcium imaging was performed across 16 different fields of view (FOV) per animal for each imaging session. Each FOV was acquired for 50 trials. For imaging fosGFP expression, a dual channel high resolution 3D stacks of RCamp1.07 and GFP were taken through the calcium-imaged area using a Ti:Sapphire laser tuned to 860 nm (Chameleon Ultra Laser; Coherent) and resolved using 630/69nm and 520/60nm emission filters (BrightLine), respectively. Following the conclusion of all experiments, animals were anesthetized and a high resolution 3D image volume was taken through the imaged area(s) for later registration with *ex vivo* tissue.

### Whisker tracking and analysis

High-speed videography of whisker movement was acquired at 500 Hz as previously described^26,54^. For analysis, whiskers were automatically traced^54^. The angle, curvature (κ), and location of the whisker tip at each time point was extracted for all traced whiskers. Using the mean whisker angle, a Hilbert transform was applied to determine whisking amplitude, phase, setpoint, and the reconstructed whisker angle (angle_RECONSTRUCTION_)^55^. The position of the rotor was automatically tracked in the video using custom scripts (MATLAB). Whisker-rotor touch was scored as events in which the tip of at least one whisker came into within <5 pixel radius of the rotor face with ‘touch onset’ defined as the first possible whisker contact at the beginning of the sample and test period and ‘touch offset’ defined as the last possible whisker contact at the end of the sample and test period. Time vectors of kinematic parameters were down-sampled to the imaging frame rate for further analysis. Behavioral sessions with sub-optimal imaging conditions in which whisker tracking failed in >20% of trials were excluded from analysis.

### *In vivo* image analysis

All image processing was performed in MATLAB as described^26,55^. Two-photon images were first motion corrected using a piece-wise rigid motion correction algorithm^56^. For behavior experiments, regions of interest (ROIs) were extracted using constrained non-negative matrix factorization and were labeled as active neurons. Remaining neurons in the FOV were manually segmented and labeled as inactive neurons. For bilateral whisker deprivation experiments, neurons across imaging sessions were identified by first registering each session image into a global reference image using ImageJ and MATLAB. For each neuron, ROIs were manually segmented in the reference image and applied to each session image for extraction of calcium signals. For analysis of fosGFP expression, 3D volume stacks across imaging sessions were aligned by landmark-based 3D affine registration using custom software (Neurotator). Automatically segmented 3D ROIs were obtained from registered *ex vivo* tissue slices corresponding to *in vivo* imaged areas and were overlaid for each imaged neuron. The mean fluorescence intensity for GFP and RCamp1.07 was extracted for each ROI. To control for variability in imaging conditions and background fluorescence, relative GFP intensity was expressed as ΔR/R_0_ = *(R*_*ROI*_-*R*_*neuropil*_)*/R*_*neuropil*_ where *R*_*ROI*_ is the ratio between GFP and RCaMP1.07 fluorescence in the ROI and *R_ROI_* is the ratio between GFP and RCaMP1.07 fluorescence in the surrounding neuropil. FosGFP neurons with ΔR/R_0_ > 0.2 were identified as high expressing.

### Spike estimation

Calcium signals were deconvolved using an Online Active Set method to Infer Spikes (OASIS), a generalization of the pool adjacent violators algorithm (PAVA) for isotonic regression (see also **Supplementary Note S.2**). First, calcium signals below baseline fluorescence (bottom 10^th^ percentile of signal intensity) were thresholded. For each cell, a convolution kernel with exponential rise and decay time constants were determined using an autoregressive model. The convolution kernel was applied to the calcium signals to obtain an initial spike estimate (*ŝ*) which was then thresholded (*ŝ* = 1) into binary events representing an estimated spike. For measurement of photon shot noise, signal-to-noise (*v)* was calculated as:

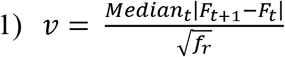

where the median absolute difference between two subsequent time points of the fluorescence trace, *F*, is divided by the square root of the frame rate, *f*_*r*_ ^57^.

### Trans-monosynaptic rabies tracing

For input mapping experiments, trans-monosynaptic rabies tracing was performed on transgenic Sst-IRES-Cre or VIP-IRES-Cre mice. AAV1-hSyn-DIO-TVA66T-dTom-CVS-N2cG virus (titer = 5.46×10^12^ vg/mL) was injected into L2/3 of S1 in the left hemisphere (1.1 mm posterior to bregma, 3.3 mm lateral, 300 μm deep) via iontophoresis at 5uA for 5 min. Four weeks later, 500 nL of EnvA-CVS-n2c(dG)-histone-EGFP virus (titer = 5×10^9^ vg/mL) was injected into S1 using the coordinates above via nanoinject. Animals were sacrificed after 7 days.

### Retrograde tracing

Retrograde tracing was performed to identify long-range axon projection targets of different molecular types of excitatory cells. Cholera toxin subunit-b (conjugated to AlexaFluor 555 (CTB-555; 500nl, 1 mg/ml; ThermoFisher Scientific) was injected into one of two major projection targets of S1 (*n* = 6 mice for each projection target): secondary somatosensory cortex (0.7 mm posterior to bregma, 4.2 mm lateral, 300-500 μm deep), or primary motor cortex (1.1 mm anterior to bregma, 0.6 mm lateral, 300-500 μm. Seven days after injection, brains were perfused and fixed in 4% PFA. Coronal tissue section (150 μm) were cleared and subjected to HCR-FISH. Multi-channel image stacks were acquired in L2/3 of S1 with FISH probes in 488 and 647 and CTB labelling in 561 using LSM confocal microscope. Percentage of CTB positive cells expressing each gene was manually quantified in ten randomly selected 150×150 μm^2^ subregions of L2/3 (ImageJ).

### *Ex vivo* tissue preparation

At the conclusion of *in vivo* imaging experiments, the cranial window was removed. Several small punctures were made in tissue surround the *in vivo* imaged area using a glass pipette dipped in lipophilic dye (SP-DiIC_18_(3) (1,1’-Dioctadecyl-6,6’-Di(4-Sulfophenyl)-3,3,3’,3’-Tetramethylindocarbocyanine; Fisher Scientific, cat no. D7777). Punctures were made in an asymmetrical pattern such that the orientation of the slice could be determined based on their locations. A stereoscopic image was taken of the punctures relative to blood vessels. The animal was then euthanized and transcardially-perfused using 1X PBS followed by 4% paraformaldehyde (PFA; 32% stock (wt/vol); Microscopy Sciences, cat. no. 15710-S). The brain was removed and further fixed in 4% PFA overnight at 4°C. The following day, the brain was mounted in 1.5% agarose gel (Agarose Molecular Bio Grade (100g); IBI Scientific, cat. no. IB70040) and sliced tangentially, parallel to the imaging plane in 150-300 um sections using a vibratome (Leica VT1000S Vibratome; Leica Biosystems). Slices were cleared using PACT-CLARITY procedure previously described^14,19^ For input mapping experiments, three or four 150 μm-thick coronal slices from each animal centered around the injection site were selected for further processing.

### Probe set and barcode design

HCR-FISHv3.0 probe sets consist of a target sequence that binds to mRNAs of interest paired with one of three orthogonal HCR hairpin amplifiers (B1, B2, B3) conjugated to either AlexaFluor488, AlexaFluor546, AlexaFluor647, AlexaFluor750, AlexaFluor790^20^ (Molecular Instruments, Inc.). Target binding sequences for transcripts were determined from sequences deposited in NCBI RefSeq (www.ncbi.nlm.nih.gov/refseq/) with the exception of RCaMP1.07 obtained from^21^. Order and lot numbers for all probe sets are listed (**Supplementary Table 1**).

For behavior experiments, a barcode scheme was implemented for gene readout based on a Hamming code similar to as previously described^12^. The barcode contained two readout channels (B1-647 and B2-488). Readout of one gene was assigned to only one channel and encoded in two out of five rounds of staining. For error-robust encoding, a Hamming distance of 2 was used such that at least 2 errors were required to switch from one readout to another. For cellular-resolution readout, genes were assigned to readouts such that no co-expressed genes were present within one round and channel of staining. A sixth round of staining was added to confirm the identity of excitatory neurons using *Slc17a7* in B2-488 and inhibitory neurons using *Gad2* in B1-647. For registration and alignment of neurons across *in vivo* images and multiple round of HCR-FISH, probe sets for RCaMP1.07 in B3-546 was used across all staining rounds.

For input mapping and bilateral whisker deprivation experiments, fewer genes were read out and *Fos* mRNA was quantified in fosGFP animals. For these reasons, sequential multiplexed HCR-FISH of non-overlapping probes was performed instead of using overlapping probes as in the barcode scheme. Since GFP was expressed in both sets of tissue, readout of gene expression was used performed with B2-647 and a mixture of B1-750:B1-790 (1:1). Genes identifying cell types were selected based on results from single cell RNA sequencing of mouse S1 neurons using SMART-Seq v4 and 10x Genomics Chromium platform as previously described^22^ (portal.brain-map.org/atlases-and-data/rnaseq).

### Multiplexed hybridization chain reaction fluorescent *in situ* hybridization

HCR-FISH was performed with modifications to the v3.0 protocol^13^. Fixed, cleared samples were incubated in 500 μL of 30% probe hybridization buffer (30% formamide, 5x sodium chloride sodium citrate (SSC), 9 mM citric acid,0.1% Tween 20, 50 μg/mL heparin, 1x Denhardt’s solution, and 10% low MW dextran sulfate) at 37°C for 30 minutes. Samples were then moved to probe solution overnight, which comprised of 500ul of 30% probe hybridization buffer and 2 μL of initiator probes (1 μL of odd probe and 1 μL of even probe, taken from 2 μM probe stock solutions) at 37°C. The next day, samples were washed four times at 37°C for 30 min in 500 uL of the following solutions, respectively: 75% probe wash buffer + 25% 5x SSC (Sigma Aldrich), 50% probe was buffer + 50% 5x SSC, 25% probe wash buffer + 75% 5x SSC, and 100% 5x SSC. Samples were then moved to 500 uL of amplification buffer (5x SSC + 0.1% Tween 20 + 10% low MW dextran sulfate) at RT on a shaker for 30 minutes. 10 μL of 3 μM stock hairpin amplifier solution was snap cooled by heating to 95°C for 90 seconds and then allowed to cool in a dark drawer for 30 min. 10μL of both snap-cooled odd and even hairpins were added to 500 μL of amplification buffer to form the final amplification solution. Samples were moved to final amplification solution and allowed to amplify at RT overnight. The following day, samples were removed from the amplification solution and washed in 5x SSC three times at RT: 30 minutes, 30 minutes, and 15 minutes. Between staining rounds, *in situ* probes were removed using DNAse (Dnase I recombinant; Sigma Aldrich ct. no. 04716728001) as described^15^. Samples were incubated for 30 min at RT in 500 μL of 1X incubation buffer (40 mM Tris-HCl, 10 mM NaCl, 6 mM MgCl2, 1 mM CaCl2). Samples were then moved to 500 μL of 1X incubation buffer with Recombinant DNase I (10 U/μL) for 4 hours at RT. Samples were then washed three times for 30 minutes at RT in 500 μL of MT Probe Wash Buffer. For bilateral whisker deprivation experiments, cell nuclei were stained using DAPI during the first round of HCR-FISH staining (DAPI Fluoromount-G; SouthernBiotech).

### Confocal imaging

Tissue sections were mounted on 75mm × 25 mm glass microscope slides (Fisher Scientific) with Fluoromount-G (SouthernBiotech) and 50mm × 22mm cover glass (Fisher Scientific). Images were acquired on one of two confocal systems: 1) a Nikon C2+ Si spectral laser scanning microscope (LSM) with Nikon Plan Apo λ 20x/0.8NA, air objective, and 0.4094×0.4094×1 μm^3^ XYZ image voxel size, or 2) a Nikon Ti2-E body with Yokogawa Spinning Disk and Nikon CFI Apo LWD 40×/ 1.15NA, water immersion objective, and 0.1625×0.1625×0.4 μm^3^ XYZ image voxel size. Only one system was used across multiple imaging rounds for a given tissue sample. For LSM-acquired images, a single image volume was sufficient to cover the *in vivo* imaging region. For spinning-disk-acquired images, multiple image tiles spanning the *in vivo* imaged region were acquired and later assembled using TeraStitcher^58^. For ‘post-cleared’ tissue prior to HCR-FISH staining, SP-DiIC_18_ (3) and endogenous RCaMP1.07 expression was acquired using 488 and 561 imaging channel, respectively. For HCR-FISH stained tissues, 488, 561, 647, or 785 imaging channels used to visualize readout hairpins. DAPI staining was acquired using the 405 imaging channel.

### *Ex vivo* image analysis

In order to re-identify neurons imaged *in vivo* and across multiple rounds of HCR-FISH, all acquired images were registered to a common reference image volume. First, 2D frame-averaged structural images taken from the behavioral session were registered into an *in vivo* two-photon 3D image stack using landmark-based 2D affine transformation (MATLAB). Next, one round of HCR-FISH staining was designated as the common reference image volume. The *in vivo* volume including registered behavior sessions, ‘post-cleared’ volume, and all other HCR-FISH volumes were then registered to the reference image volume based on endogenous protein or HCR-FISH stained mRNA RCaMP1.07 expression. Landmark-based 3D thinplate registration was performed to generate a coarse alignment of the image volumes using custom software (Neurotator). This coarse alignment produced a registration accuracy of <5μm. For HCR-FISH stained volumes, thinplate registration was performed on the 561 imaging channel containing RCaMP1.07 expression was then applied to the remaining image channels. (**Supplementary Fig. 1**). For input mapping experiments, HCR-FISH stained volume registration was performed on the 488 imaging channel containing GFP expression of input neurons.

Following coarse alignment, an automatic fine-scale 2D rigid alignment was applied to each z-frame of HCR-FISH stained volumes using matrix-multiply discrete Fourier transform^59^. Registration was performed on the 561 imaging channel for behavior and bilateral whisker deprivation experiments or the 488 channel for input mapping experiments and then applied to the remaining image channels. For LSM-acquired images, fine-scale alignment was performed on the entire image stack. For spinning-disk-acquired images, sub-volumes corresponding to regions containing individual neurons imaged *in vivo* were first isolated and then subjected to fine-scale alignment in order to reduce CPU processing time.

In order to characterize gene expression in identified neurons, 3D cell segmentation was performed on image volumes. For behavior experiments, segmentation was performed using RCaMP1.07 expression. Following fine alignment, image stacks of RCaMP1.07 transcript expression across all imaging rounds were merged into a single volume stack. Segmentation was performed on the merged volume stack to obtain a single 3D segmented *ex vivo* ROI applied to each imaging round. For LSM-acquired images, segmentation was performed on HCR-FISH stained somatic RCaMP1.07 using Modular Interactive Nuclear Segmentation^60^. For spinning-disk-acquired images, images stacks were pre-filtered to normalize signal intensity and to label nuclei within RCaMP1.07-expressing neurons using 3D Weka Segmentation^61^. The resulting filtered images were merged with the original images and then segmented using Baxter Algorithm^62^. For whisker deprivation experiments, segmentation was performed on DAPI stained nuclei. The resulting ROI was expanded radially by 10 voxels to better encompass the somatic region. RCaMP1.07 fluorescence within each ROI was use determined to determine whether identified cells expressed RCaMP1.07. For input mapping experiments, segmentation was performed on nGFP expression. The resulting ROI was expanded radially by 10 voxels to better encompass the somatic region. All *ex vivo* segmented neurons were manually validated using Neurotator.

Segmented *in vivo* ROIs were matched with segmented *ex vivo* ROIs by a combination of point cloud registration of ROI centroids and pixel overlap. Candidate matches were identified by nearest neighbor sorting (MATLAB). Overlap was calculated as:

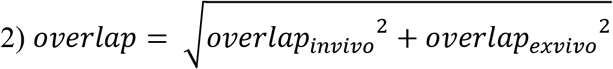

where *overlap*_*invivo*_ is the fraction of *in vivo* ROI pixels overlapping with the *ex vivo* ROI and *overlap*_*exvivo*_ is the fraction of *ex vivo* ROI pixels overlapping with the *in vivo* ROI. Candidate matches were rejected if *overlap* < 0.5. Matching ROIs were manually validated using Neurotator. *In vivo* imaged ROIs determined not be contained within the *ex vivo* tissue volume were assigned to be unlabelled (UNL) neurons and excluded from cell type identification.

For behavior experiments, binary readout of gene expression in each HCR-FISH imaging round was manually scored by visual inspection using Neurotator. For decoding, barcodes were designed to avoid overlapping gene expression for each round of staining. However, cells that co-express genes assigned to the same imaging channel can occasionally produce ambiguity in decoding. To resolve these situations, a combinatorial list of candidate barcodes was listed based on the barcode scheme used in the experiment and all possible gene expression patterns. From this list, non-unique barcodes were excluded as well as barcodes in which exhibitory and inhibitory markers were co-expressed. Co-expressed genes were identified from this plausible list of barcodes. Any cells with ambiguous barcodes were assigned as UNL neurons and excluded from cell type identification. While the use of a Hamming code enables error correction of the barcode readout, no error correction was performed.

For input mapping and bilateral whisker deprivation experiment, the Python library *starFISH* was implemented for single mRNA spot detection and quantification^63^. Spots were detected using *min_sigma* = 0.5, *max_sigma* = 7, and *num_sigma* = 15. Spot detection threshold was set using Otsu’s method from the *scikit-image* library. Spot detection performance was verified visually. The number of spots were quantified within each ROI and expression levels were expressed as the number of spots per μm^3^. For each round of staining, background spot density was quantified from the surrounding neuropil across tissue subvolumes and binary readout was determined if the ROI spot density exceeded 2 times the 95^th^ percentile of background spot density.

Cell types were identified using gene expression patterns. While inhibitory classes and subclasses are identified by largely non-overlapping genes, excitatory cell types were identified by combinatorial expression patterns. Specifically, Adamts2 neurons were identified by co-expression of *Fst* and *Ngb*. Ba1za neurons were defined by expression of either Penk or Rrad and co-expression of either *Fst*, *Ngb*, *S100a6*, or *Coch*. Agmat neurons were identified by expression of either *S100a6* or *Coch* but no co-expression of *Fst, Ngb*, *Rrad*, or *Penk*. Neurons not conforming to these expression patterns were assigned as UNL neurons.

For input mapping experiments, the density of input neurons along L2/3 was calculated as the number of nGFP+ cells within a 50μm sliding window along laminar depth divided by the total number of nGFP+ cells within L2/3 for each imaged coronal slice.

### Analysis of stimulus- and behavior-related activity

To assess the link between neuronal responses and various explanatory variables, a Poisson GLM was fit to each neuron’s deconvolved spike train across a recording session^27^ (see also **Supplementary Note S.3**). The model defines probability in terms of a time-varying spike rate, λ_*t*_, given by:

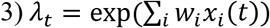

where *x*_*i*_*(t)* represents the time course for the *i*th explanatory variable, and *wi* represents the effect of this variable on the neuron’s probability of spiking^64^. The regularized log-likelihood of a spike train for an individual trial, assuming a Poisson distribution, is given by:

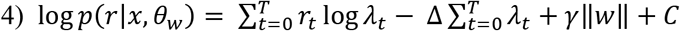

where Δ is the time bin size, T is the number of time bins in the trial, *r*_*t*_ is the spike count at time *t,* and γ is the lasso penalty term which encourages sparseness on the weights, *w.* All GLMs were fit using MATLAB’s lassoglm function with a Poisson link function, 6 penalty values (γ), and 4 fold cross-validation.

To represent the time-course of task variables *xi(t)*, boxcars were constructed which were designated as “true/1” during time-periods of interest and “false/0” elsewhere. Since task timing was fixed across trials, each trial could be broken up into six time-periods of fixed start time and duration: “pre”, “sample”, “early delay”, “late delay”, “test”, and “report”. Complementary covariates were represented as independent boxcars (ie. anterior stimulus and posterior stimulus; match and non-match; lick and no lick) to capture any asymmetry in the neuron’s response. To assess if a neuron’s activity could be explained by trial information occurring at multiple points in the trial, some task variables were represented in multiple time periods. For a given trial, the sample stimulus direction was represented as three boxcars: during the sample stimulus presentation period, during the early delay period, and during the late period. Trial category and choice were represented as a boxcar during both the test period and the report period.

Whisker kinematics and population activity were represented as continuous variables across the entire trial. All whisker kinematic features were z-scored and down-sampled to the imaging rate before including as a covariate. Whisker touch onset and offset was represented as a boxcar of width 5 bins (150 ms) centered at the touch event. The coupling covariate was calculated for each neuron by excluding the given neuron’s spike train, then calculating the non-negative matrix factorization (NMF) of the matrix of spike trains of all other neurons in the session^28^. A NMF of rank = 1 used when assessing encoding of task factors. For assessing coupling across varying NMF ranks, each rank was included as a covariate in the model and the coupling factor included all covariate ranks.

Related covariates were grouped together into ‘task factors.’ For each task factor, a partial model was constructed that excluded the covariates associated with this task factor. Any increase in deviance from the full model to the partial model therefore resulted from the exclusion of this task factor’s covariates. Akaike Information Criterion (AIC) was used to compare deviance between partial models in which different number of covariates were excluded such that:

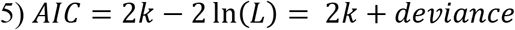

where *k* is the number of model parameters, deviance = *-2*ln*(L),* and *L* is the model likelihood. The difference in AIC (ΔAIC) between the full and partial model was calculated as:

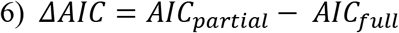

For analysis, three GLMs were constructed. Model 1 (task GLM) was constructed to assess task-related coding of neurons using task variables and task factors shown in **Supplementary Fig. 8**. Model 2 (free whisking GLM) was constructed to assess neuronal responses to free whisking. Only whisker kinematic covariates included in Model 1 were included in this model. Task factors were calculated for each of these covariates. The model only included time points consisting of the last 1.4 s prior to sample period touch onset and the last 1.1s prior to test period touch onset, consistent with periods of no whisker-rotor contact preceding stimulus presentation.

Model 3 (cell type coupling GLM) was constructed to assess coupling of activity of each major cell subclass or type and consisted of all task variables in Model 1 with the exception of the ‘coupling” covariate for all simultaneously neurons. Instead, seven coupling covariates for the six main cell populations (Adamts2, Baz1a, Agmat, Pvalb, Sst, Vip) and all other, unlabeled cells in the session (UNL) were included and consisted of the NMF of rank = 1 of the spike trains for simultaneously recorded neurons sorted according to each cell subclass or type. Task factors in this model corresponded to each of the main cell type coupling covariates.

For bilateral whisker deprivation experiments, the mean firing rate to anterior and posterior whisker deflections were calculated. To account for direction selectivity, the maximum activity level to either stimulus direction was used as measure of a neuron’s stimulus response (SR). The change in stimulus responsiveness during BD was expressed as *(SR*_*post*_ - *SR*_*pre*_)*/(SR*_*post*_ + *SR*_*pre*_) where *SR*_*post*_ is the stimulus response at either day 1 or day 5 and *SR_pre_* is the mean stimulus response across Day −2,−1, and 0.

### Network analysis

Network analysis was performed to analyze functional connectivity between cell types across different task conditions. Task-specific networks were constructed, composed of neurons exhibiting significant ΔAIC (*P* < 0.01, χ^2^ test) to task factors in Model 1 (task GLM). A ‘non-coding’ network was composed of neurons that did not exhibit any significant coding for any the task variables. Information contained in each network was derived from results from Model 3 (cell type coupling GLM). A network contained six nodes that were each composed of neurons belonging to a given major cell population. An input node was defined as the neuron modeled in the cell type coupling GLM. An output node was defined as the cell types coupling factor explaining the activity of the modeled input neuron node. A directed network edge from the output node to the input node was defined as the ΔAIC of a particular cell type coupling factor that explains the activity of the input node, averaged across all input node neurons.

Network strength was calculated as the mean edge weight for all edge weights in the network. The input node strength was calculated as the sum of weights of inward directed edges from output node to the input node divided by the number of inward directed edges. The output node strength was calculated as the sum of weights of outward directed edges from input node to the output node divided by the number of outward directed edges. To test the strength and stability of functional connections across task conditions, the overall connection strength was calculated as the mean edge weight across all task networks. The stability of the connection was calculated as the coefficient of variation of the edge weight across all task networks.

### Statistical procedures

No statistical methods were used to predetermine sample size. The experiments performed were not randomized and the investigators were not blinded to allocation during experiments and outcome assessment. Statistical tests used are indicated in figure legends. Error bars on plots indicate standard error of the mean (s.e.m.) unless otherwise noted.

For statistical tests of task encoding, a χ^2^ test was performed to assess the significance of the GLM ΔAIC values. A Mann-Whitney *U* test was used to compare the strength of GLM ΔAIC values between cell types. The Bonferroni-Holm method was used to correct for multiple comparisons.

For statistical tests of bilateral whisker deprivation, a one-way ANOVA test with post-hoc multiple comparison test was used to assess change in stimulus response across the imaging sessions. A Student’s *t*-test was used to assess differences in stimulus response between fosGFP neurons as well as between excitatory cell types. A χ^2^ test was used to compare fraction of stable high-expressing fosGFP cells between excitatory cell types. Standard deviation of the fraction of stable high-expressing fosGFP cells for a given cell type was obtained by bootstrapping with replacement. The fraction of stable high-expressing fosGFP cells was determined from subsets (60% of total) of cells that were randomly sampled with replacement. This process was repeated 1,000 times and the standard deviation was determined from this distribution.

For statistical tests of network analysis, to determine whether a given node strength was significantly different from nodes within the same network or between networks, bootstrap sampling with replacement was performed on the neurons representing the network. The edge strengths and node strengths were recalculated from the resulting bootstrapped data set. This process was repeated 1,000 times to obtain 95% confidence intervals for significance tests. The original node strength was then compared to the bootstrapped node strength of the other nodes in the same network or the same node across networks for within- or across-network comparisons, respectively. To determine whether a given connection was stronger and more stable than chance, permutation tests were performed by shuffling cell type labels for each input node for each task network and recalculating the overall connection strength and stability. This process was repeated 1000 times to obtain a shuffled distribution. Connections that were both stronger and more stable than the 95^th^ percentile of the shuffled distribution were considered to be above chance. The Bonferroni-Holm method was used to correct for multiple comparisons.

For statistical tests of input mapping, a Student’s *t*-test was used to assess sublaminar and cell type differences in input density between Vip and Sst classes.

## Data and code availability

The data that support the findings of this study are available from the corresponding authors upon reasonable request. Any custom written code used for data acquisition or analysis in this study is available from the corresponding authors upon reasonable request.

## Notes

### Competing Interest Statement

The authors have declared no competing interest.

