## Supplementary Figures and Notes for "Dense Functional and Molecular Readout of a Circuit Hub in Sensory Cortex"

### SUPPLEMENTARY MATERIALS

**Supplementary Figures 1-17**

**Supplementary Note S.1. Sampling of Neurons with the CRACK platform.**

**Supplementary Note S.2. Comparison of Calcium Response Properties and Spike Inference across Cell Types.**

**Supplementary Note S.3. Task Encoding Across Neurons.**

**Supplementary Note S.4. Measuring Functional Connectivity using Cell Type Coupling.**

**Supplementary Table 1. Transcript sequences for HCR-FISH probe sets.** Sequences used to generate probe sets for all transcripts investigated in this study. Lot numbers from Molecular Instruments are also included.

**Supplementary Table 2. Sample sizes for *in vivo* imaged neurons.** Number of identified neuronal subclasses and cell types for each animal imaged during behavior.

**Supplementary Video 1. CRACK platform overview.** Animation depicting cortical neurons functionally characterized *in vivo* and subsequently transcriptionally profiled using multiplexed HCR-FISH.

### SUPPLEMENTARY FIGURES

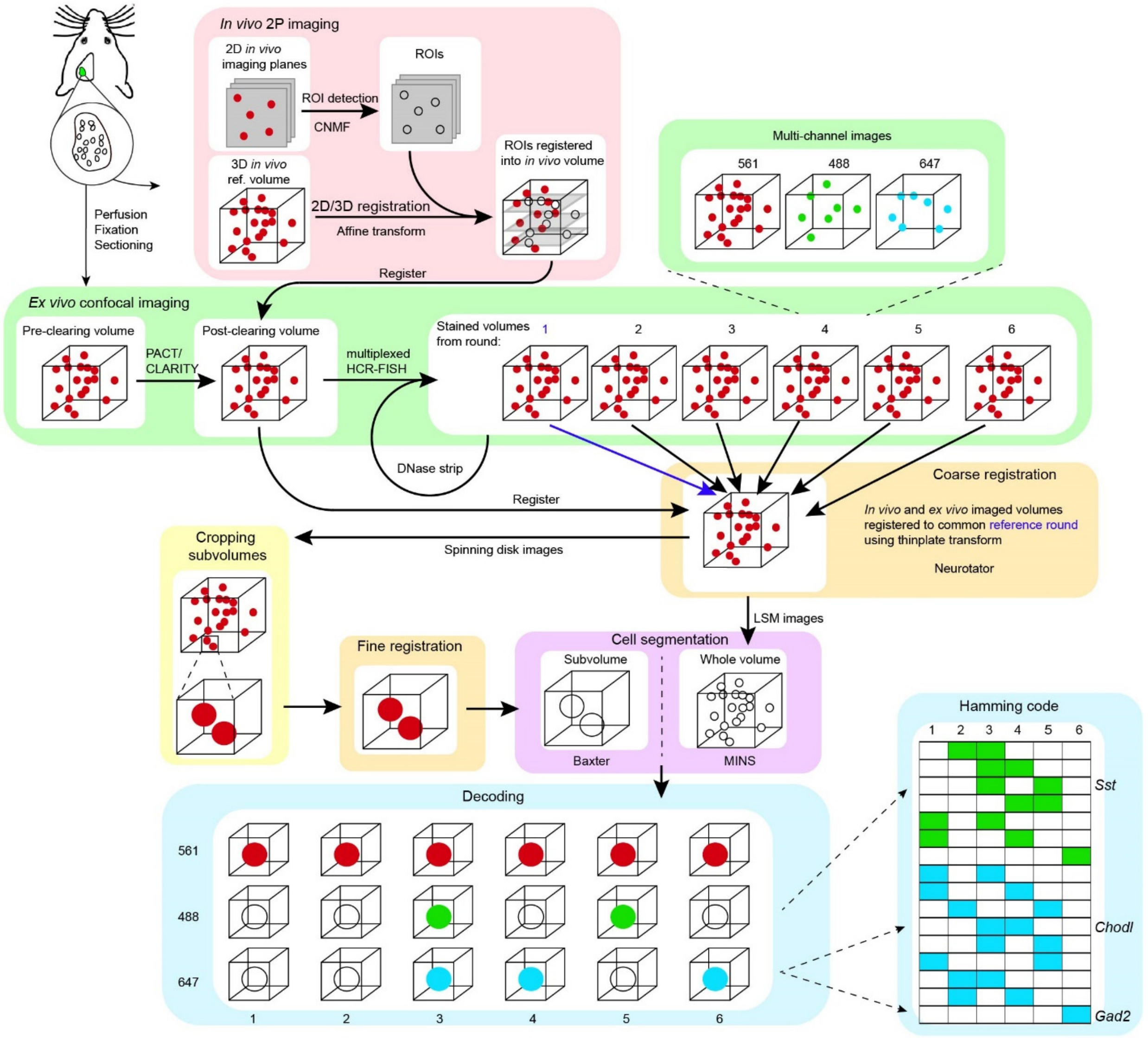


**Supplementary Figure 1. CRACK platform workflow.** *In vivo* two-photon calcium imaging of neuronal activity was performed during behavior. Two-dimensional regions of interest (ROIs) corresponding to active neurons were segmented and registered into an *in vivo* 3D reference stack. Following tissue extraction and clearing, *in vivo* area was re-identified and 3D confocal image volumes were acquired prior to and following multiple rounds of HCR-FISH staining. *In vivo* and *ex vivo* image volumes were coarsely registered to a common reference volume based on RCamp1.07 expression (561 channel). For LSM confocal images, 3D cell segmentation was performed on the entire imaging volume. For spinning disk images, imaging volumes were first cropped into sub-volumes, further subjected to fine alignment, and then 3D cell segmentation. Segmented *in vivo* and *ex vivo* ROIs are matched. HCR-FISH staining in the complementary channels (488 and 647) are decoded to obtain gene expression patterns.

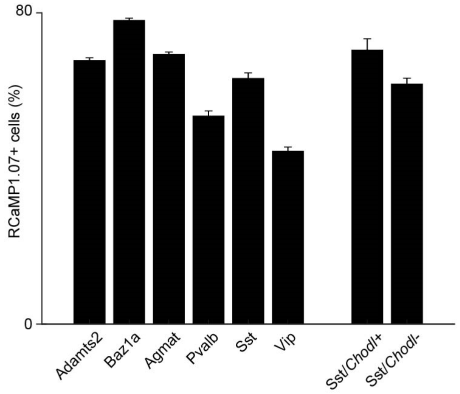


**Supplementary Figure 2. Preferential labeling of L2/3 cell types by AAV.PhP.eB vectors.** Distribution of RCaMP1.07+ neurons stained with HCR-FISH in viral injected areas across cell types. Error bars; s.d. from bootstrap analysis. $n$ = 18,506 neurons from 3 animals.

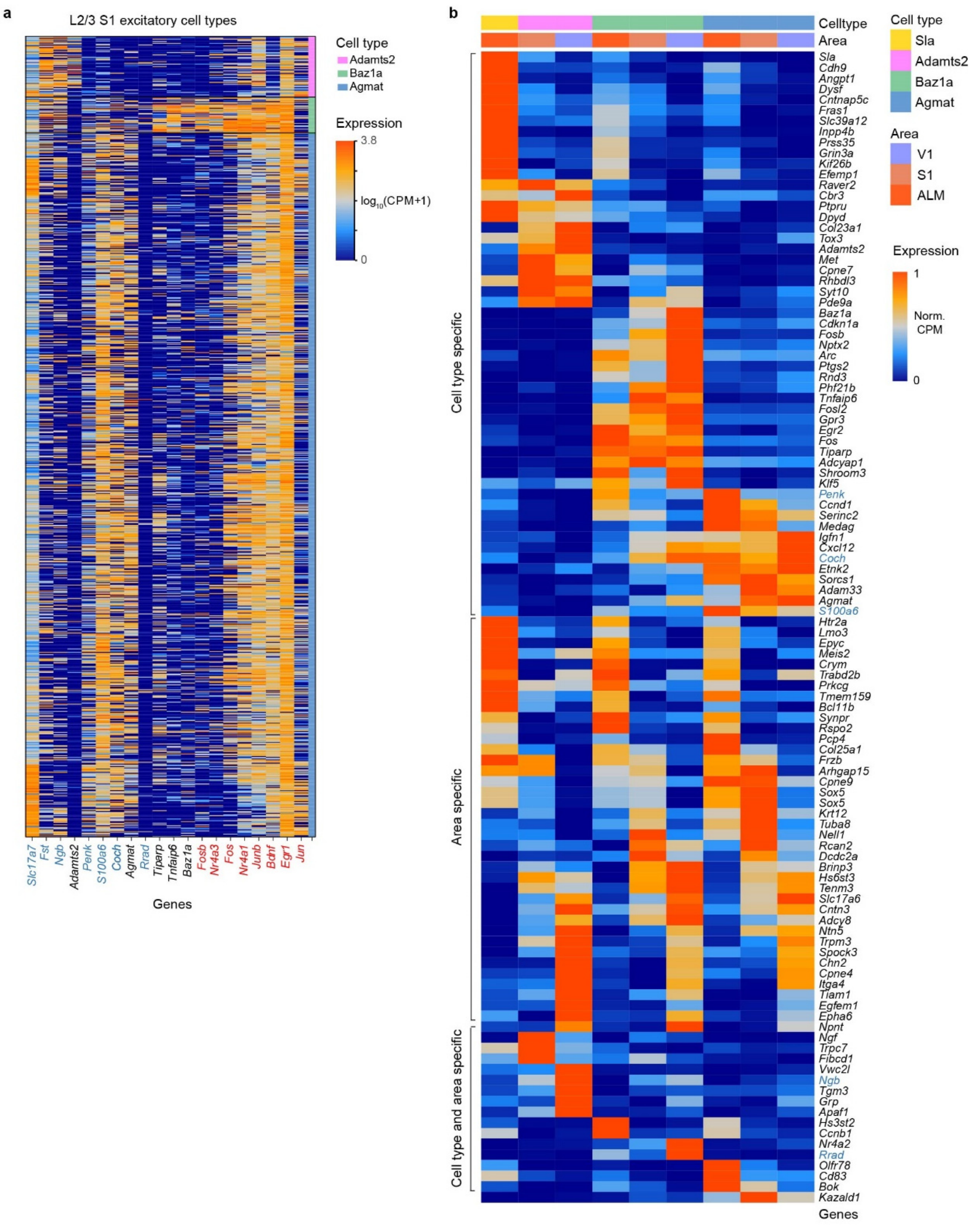


**Supplementary Figure 3. Single cell RNAseq analysis of L2/3 excitatory cell types. a,** Three major molecularly defined excitatory types in L2/3 S1 identified by single cell RNA sequencing. Expression patterns of select genes for individual cells are shown. Genes used in the CRACK platform are denoted (blue). Baz1a neurons show selective expression of immediate early genes (red). Additional selective expression of non-immediate early genes (black) indicates that Baz1a is a static excitatory cell type. **b,** Comparison of L2/3 excitatory cell types across S1, V1, and ALM. Genes with cell type specificity, area specificity, and both area and cell type specificity are shown. CPM, counts per million.

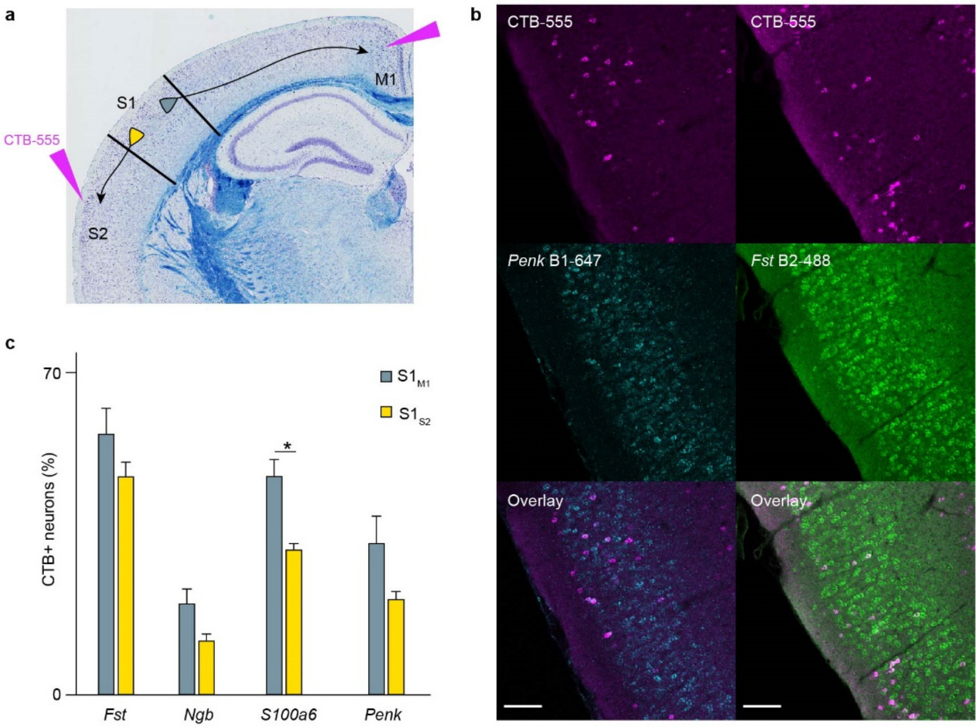


**Supplementary Figure 4. Genes defining L2/3 excitatory cell types do not label distinct populations of projection neurons.** **a,** Injection scheme for retrograde tracing experiments. Cholera Toxin subunit-b (CTB-555) was injected into one of two areas: secondary somatosensory cortex (S2) or primary motor cortex (M1). **b,** Example confocal images of HCR-FISH stained tissue section in L2/3 S1 labelling of CTB-555 from M1 (magenta), *Fst* (green), and *Penk* (cyan). **c,** Percent of identified projection neurons expressing genes defining excitatory cell types (*$P < 0.05$, two-tailed Student's *t*-test). Error bar: s.e.m. ($n = 2734$ cells from 5 animals, $S1_{M1}$; 3446 cells from 6 animals, $S1_{S2}$). Scale bars: 100 μm.

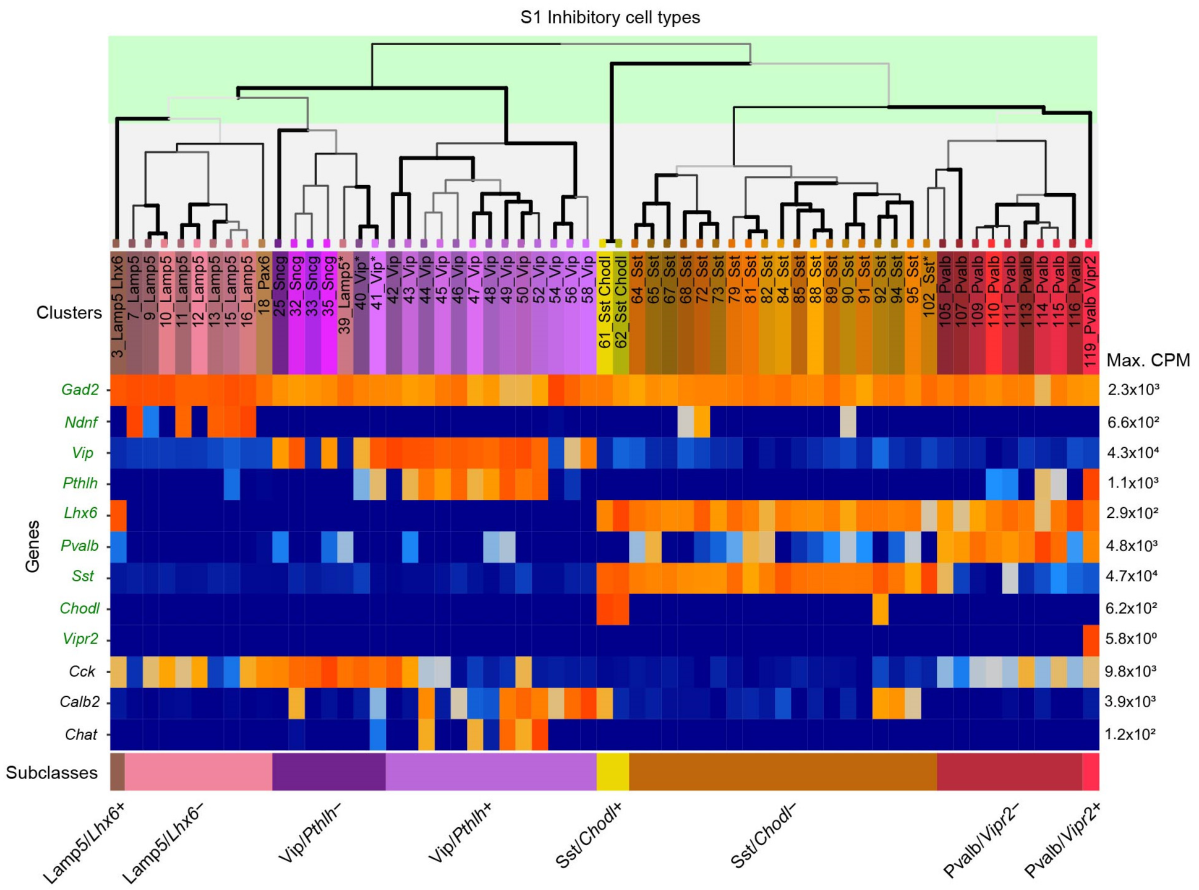


**Supplementary Figure 5. Single cell RNAseq analysis of inhibitory subclasses.** Dendrogram (top) depicts the hierarchical organization of inhibitory neurons into multiple discrete cell types. Expression profiles for a subset of genes including those selected for CRACK (green) are shown for each cluster (middle). Each gene is normalized to its maximum expression value. Selected subclasses and types for CRACK (bottom) correspond to the hierarchical level in the shaded green region of the dendrogram. CPM, counts per million.

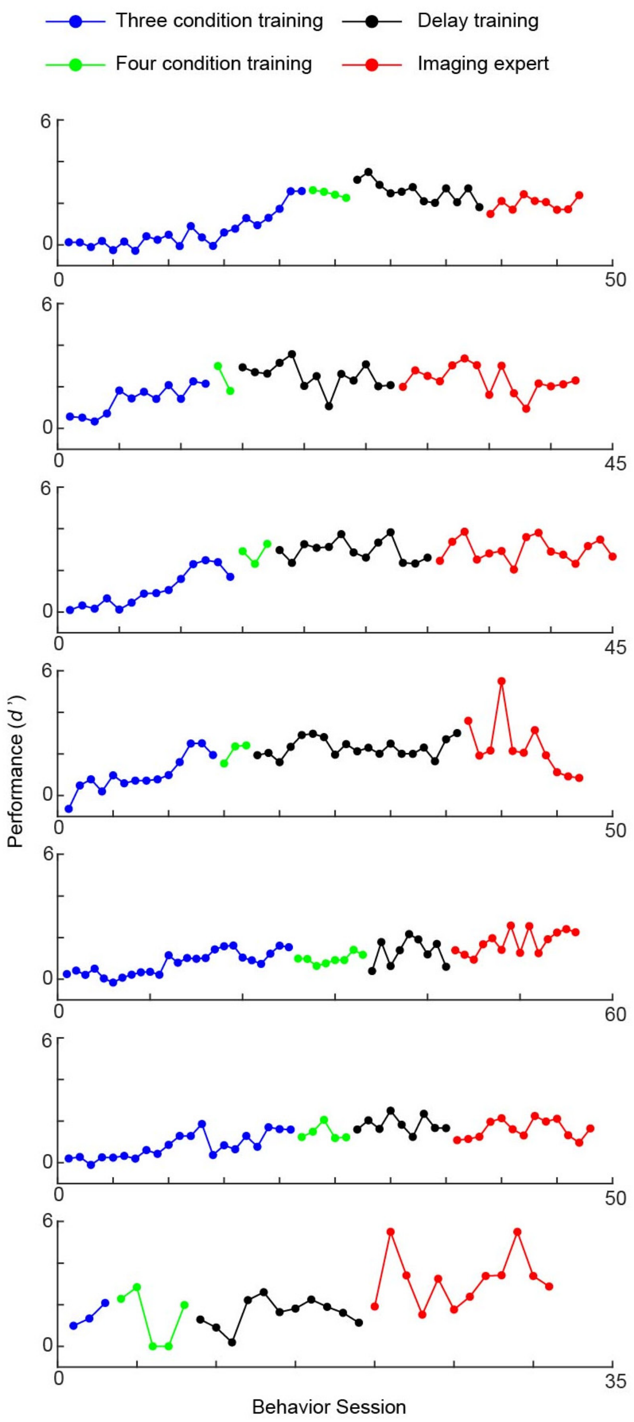


**Supplementary Figure 6. DNMS task performance during training and imaging for individual animals.**

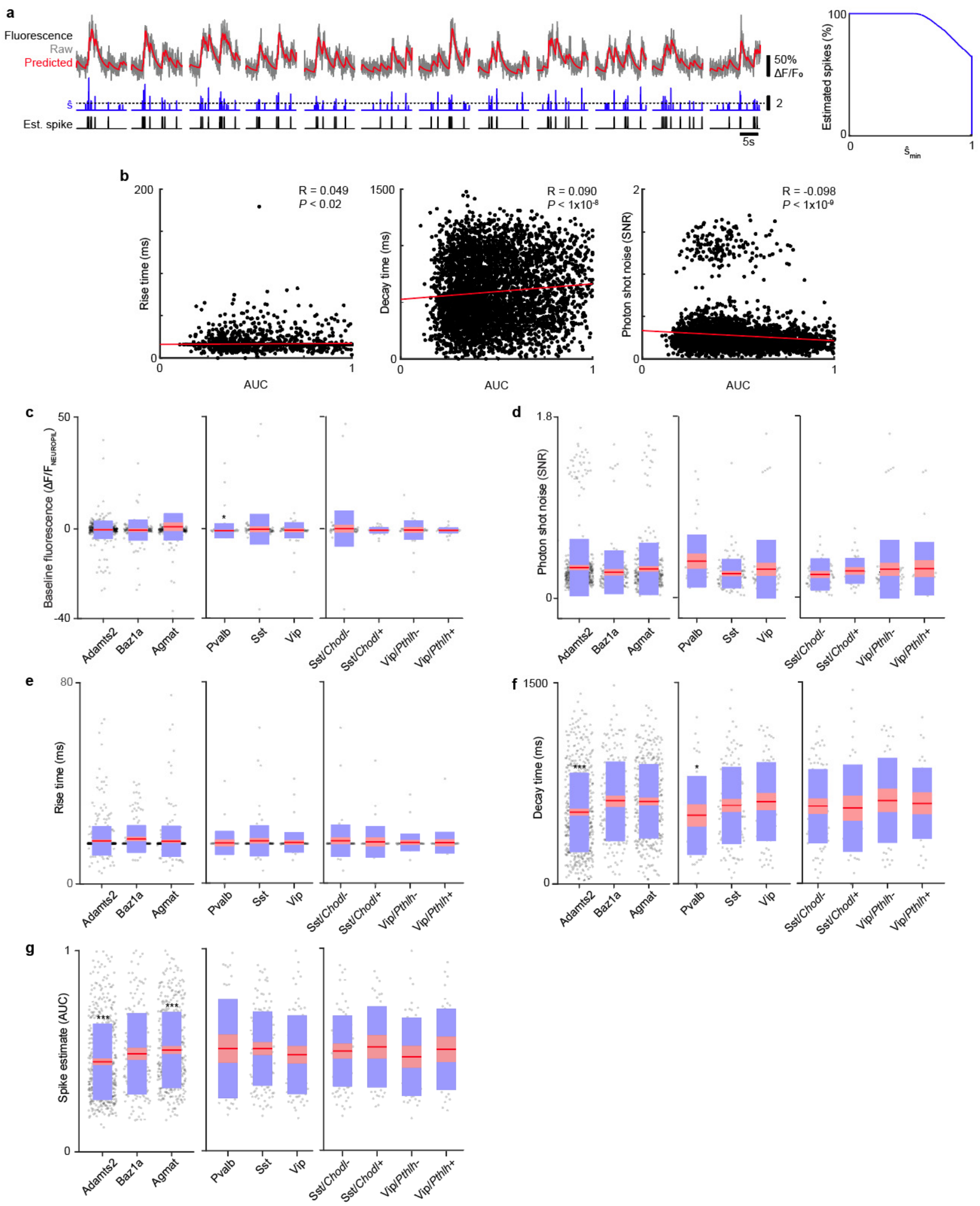


**Supplementary Figure 7. Calcium kinetics and spike estimation across cell types. a,** Example of spike estimate from calcium signal using OASIS (left). Raw signals are fit using an auto-regressive model. The predicted trace is then deconvolved into a spike estimate ($\hat{s}$) which is thresholded into binary spikes (left). Receiver operator characteristic (ROC) showing relationship between model performance across varying $\hat{s}$ threshold (right). **b,** Scatter plot showing relationship between spike estimation performance from ROC analysis vs. modeled calcium rise time (left), modeled decay time (middle), and photon shot noise (right) across individual neurons (Pearson's correlation). **c,** Baseline fluorescence intensity across cell types. **d,** Photon shot noise across cell types. **e,** Modeled rise times across cell types. **f,** Modeled decay times across cell types. **g.** Spike estimation performance across cell types. For **c-f**, red line: mean; pink region: 95% confidence interval; blue region: s.d. (* $P < 0.05$; *** $P < 1\text{x}10^{-5}$; two-tailed Student's *t*-test with Bonferroni-Holm post-hoc correction).

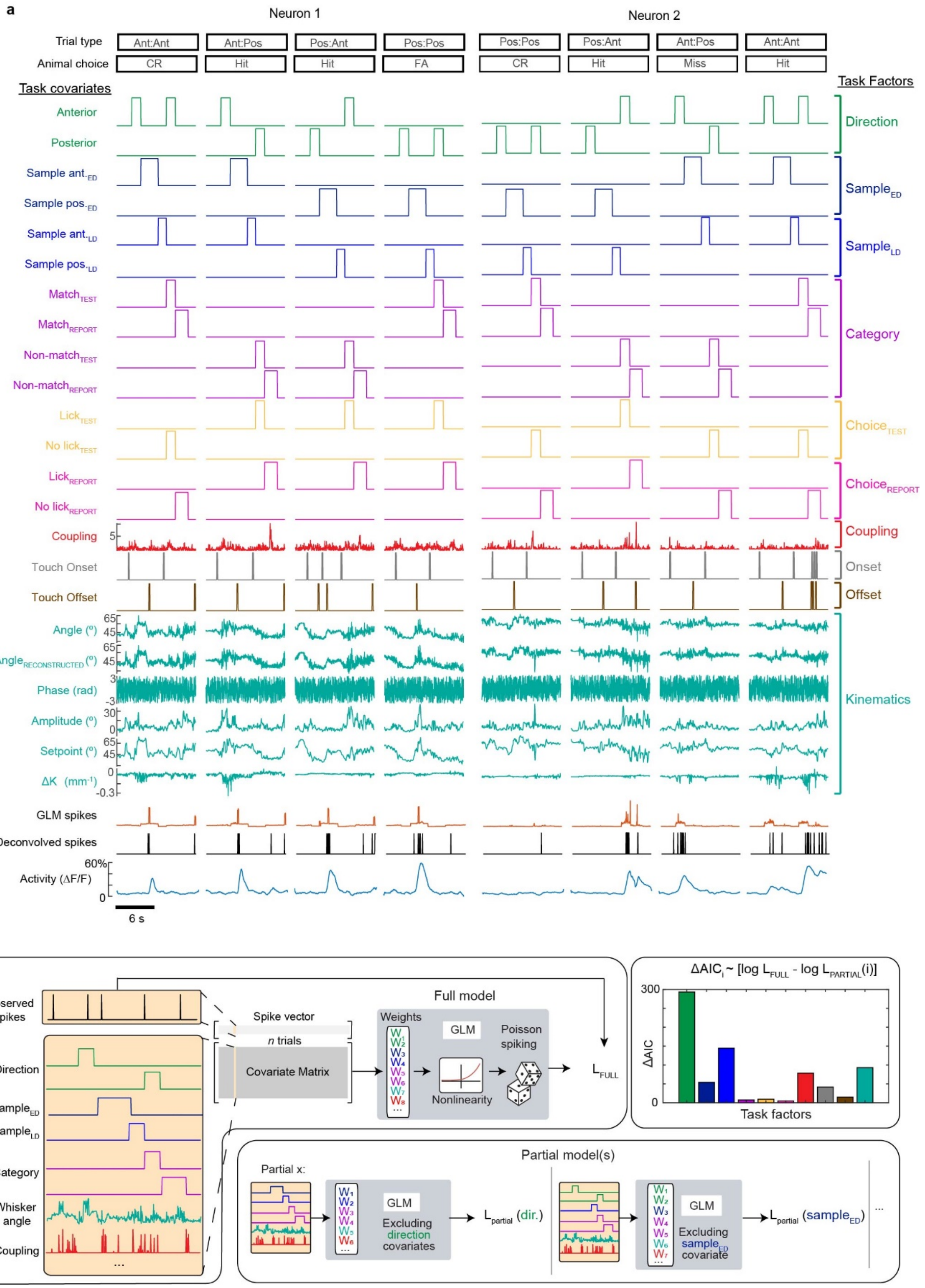


**Supplementary Figure 8. Task encoding generalized linear model. a,** Overview of covariate representations and their corresponding task factors used in in the task-related GLM for two example neurons over four trials. **b,** Schematic of full and partial models used to calculate ΔAIC for individual task factors.

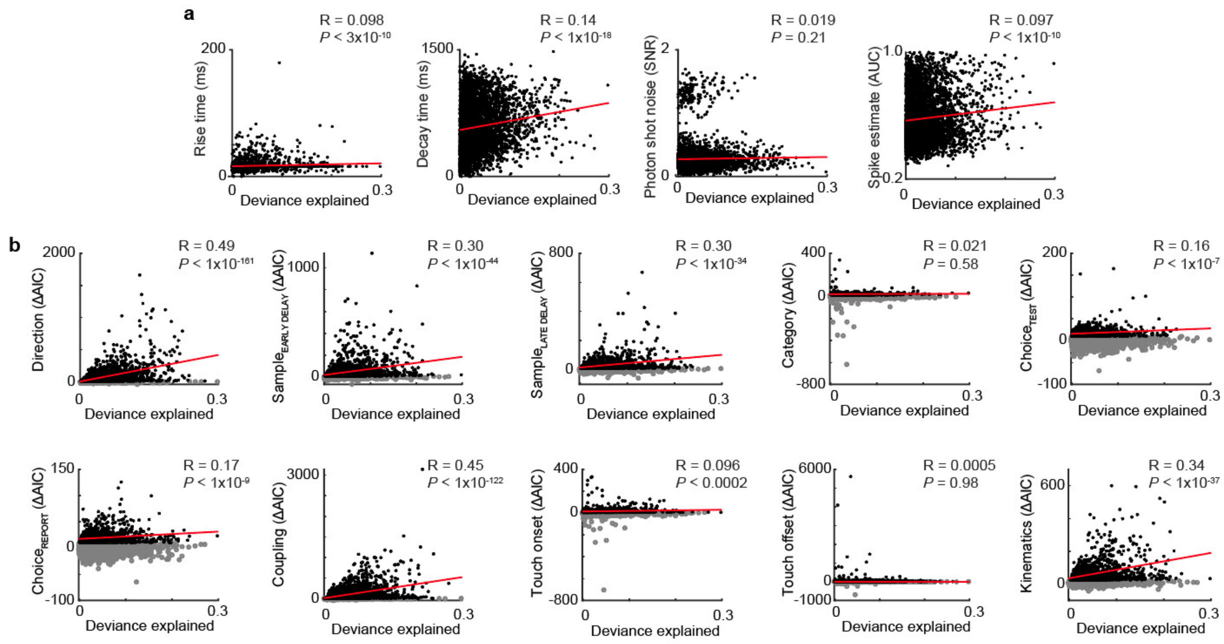


**Supplementary Figure 9. Analysis of GLM model performance. a,** Relationship between spike estimation obtained from OASIS vs. full model GLM fit for modeled rise time, modeled decay time, photon shot noise, and spike estimate performance across individual cells (Pearson's correlation). **b,** Relationship between encoding strength of task factors vs. full model GLM fit across individual cells (Pearson's correlation). Significant ($P < 0.01$; black) and non-significant (grey) ΔAIC are shown determined by $\chi^2$ test.

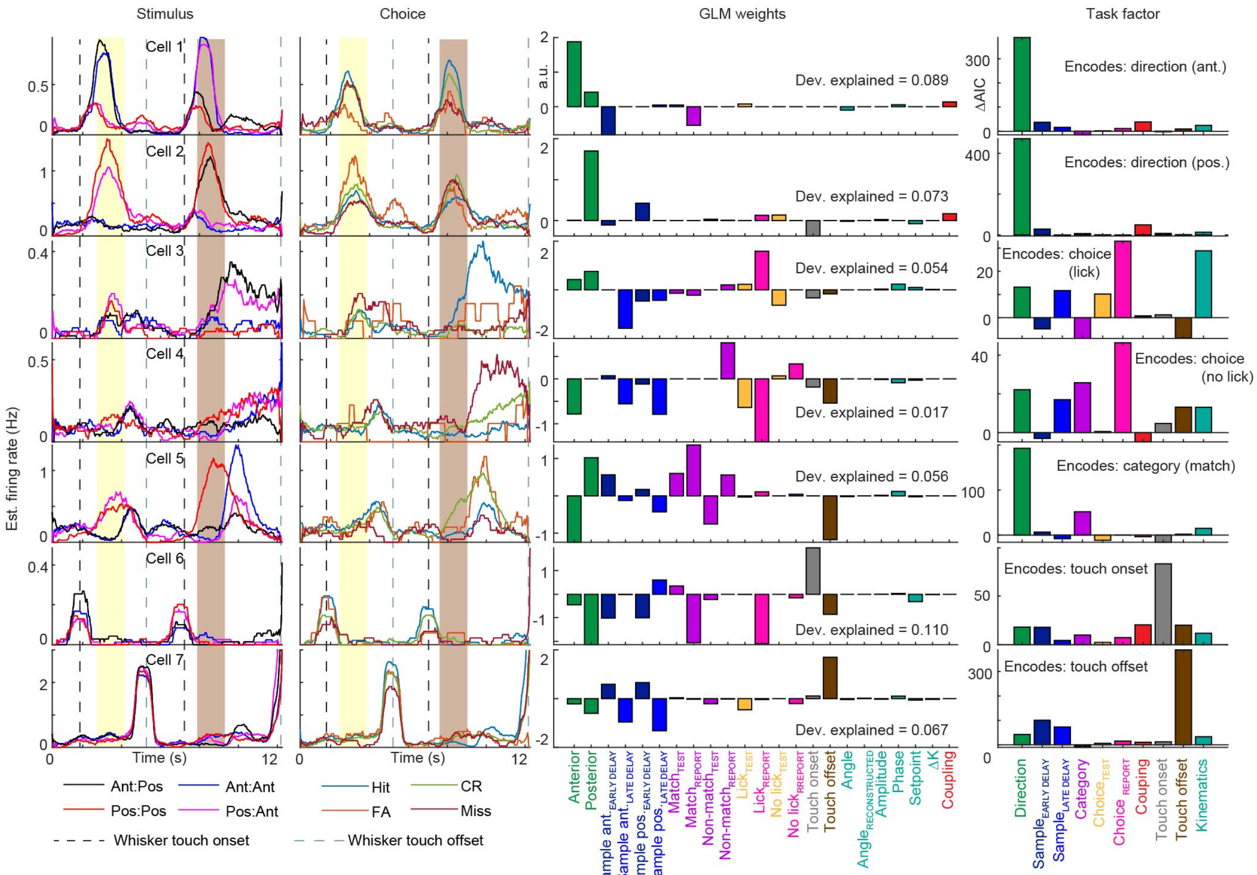


**Supplementary Figure 10. Examples of task encoding neurons.** For seven example neurons, average firing rates are shown for stimulus conditions (far left) and animal's decision (middle left). Whisker touch onset and offset are also shown. Full model GLM weights for each covariate and deviance explained are shown (middle right). Encoding strength for each task factor are shown (far right).

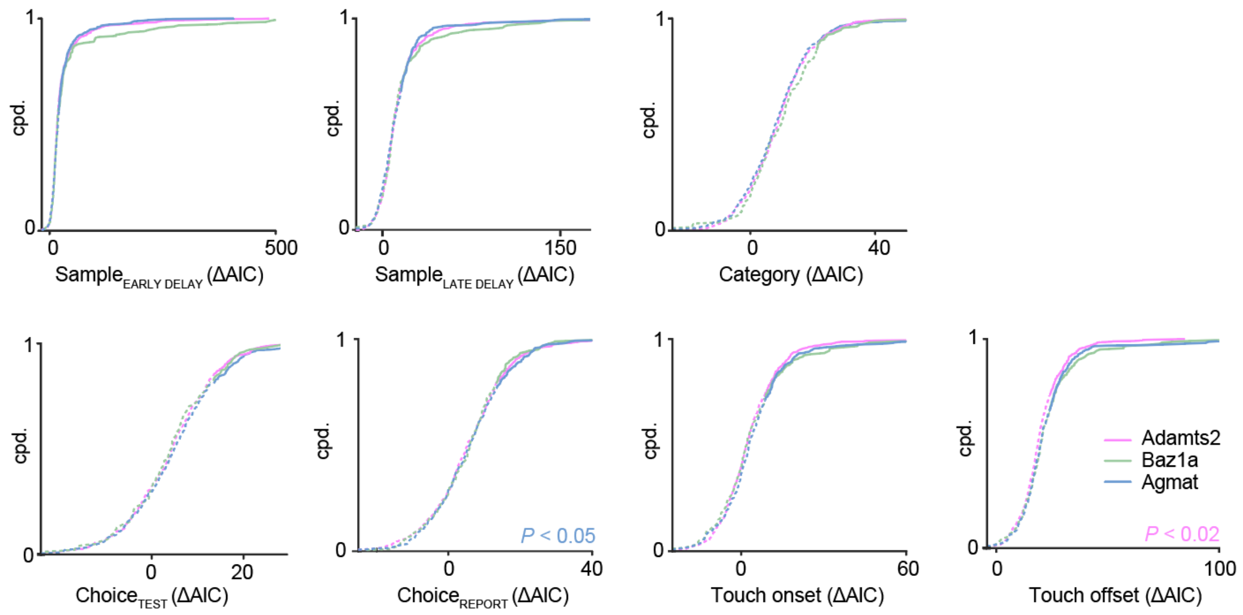


**Supplementary Figure 11. Additional task encoding across L2/3 excitatory cell types.** Encoding strength to remaining task factors shown in **Fig. 2** across excitatory cell types (Mann Whitney *U* Test). Solid and dotted lines correspond to significant ($P$ < 0.01) and non-significant encoding strengths determined via $\chi^2$ test. $n$ = 1107 neurons from 7 animals.

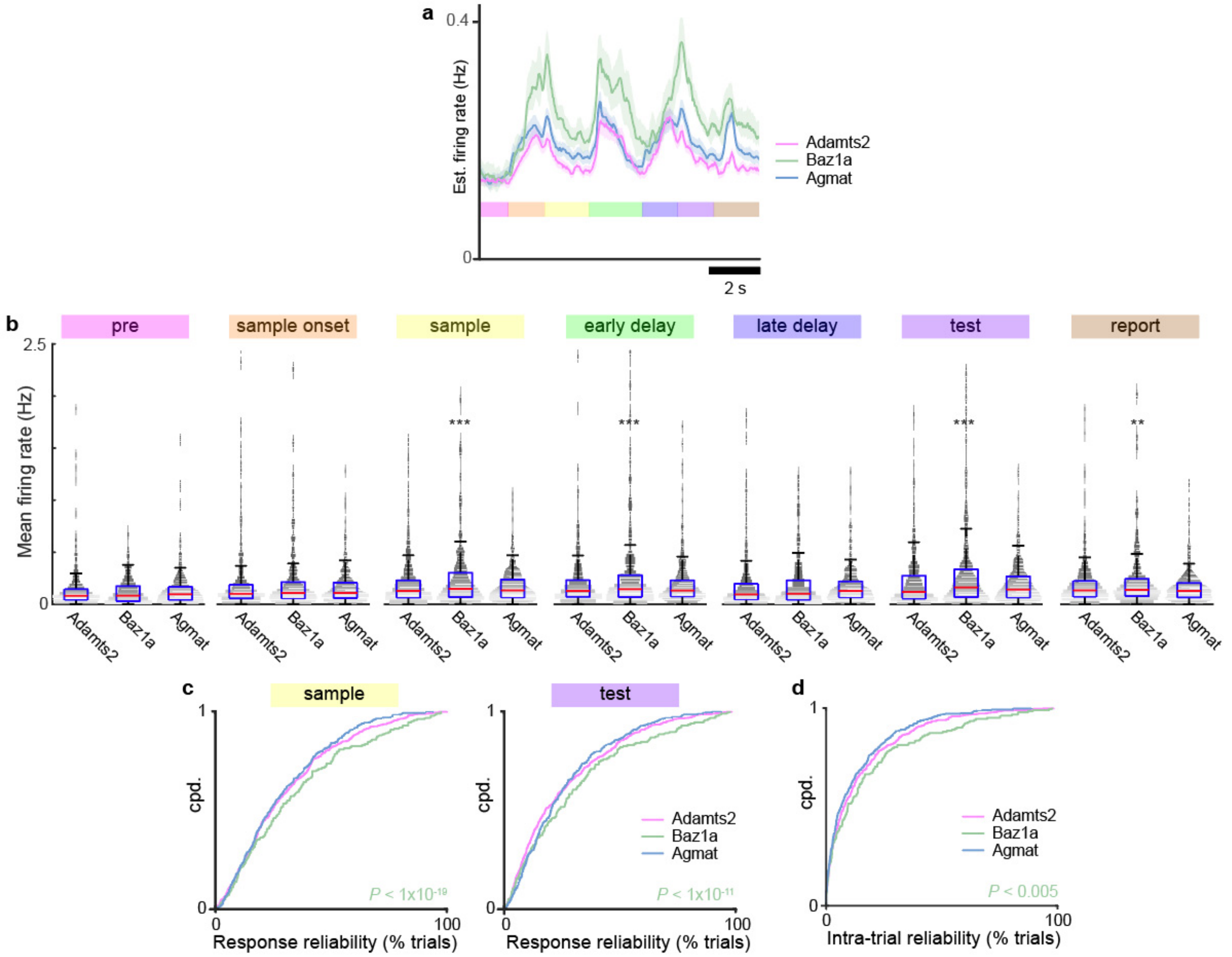


**Supplementary Figure 12. Activity levels and response reliability of excitatory cells classes.** **a,** Mean firing rate for each excitatory cell type across the trial period. Shaded regions correspond to s.e.m. **b,** Distribution of estimated firing rate responses binned across each time period for neurons belonging to the three excitatory types. Whisker box plots are overlaid over violin plots to show distribution. (** $P < 0.01$, *** $P < 0.001$, right-tailed Student's *t*-test with Bonferroni-Holm post-hoc correction). Box centerline indicates median, box limits indicate upper and lower quartiles, and whiskers indicate 1.5x interquartile range. **c, d,** Cumulative probability function of response reliability during the sample (left) and test (right) period (**c**) and intra-trial response reliability across the sample and test period (**d**) for the three excitatory types. Baz1a cells exhibit both greater response reliability and intra-trial reliability compared to other excitatory neurons (right-tailed Student's *t*-test with Bonferroni-Holm post-hoc correction). $n = 1107$ neurons from 7 animals.

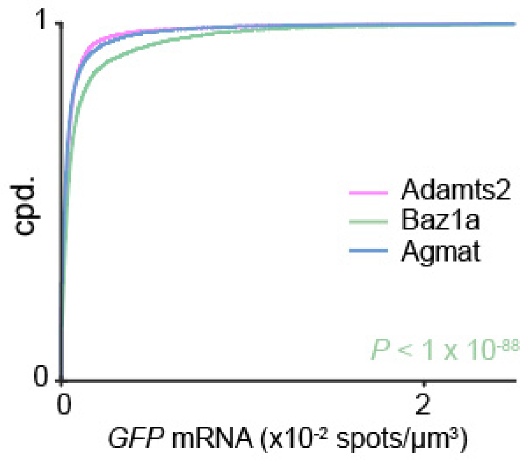


**Supplementary Figure 13. fosGFP expression across excitatory cell types.** Cumulative probability distribution *GFP* mRNA density from HCR-FISH in fosGFP mice in L2/3 excitatory cell types (Mann Whitney *U* test). $n$ = 26,396 cells from 3 animals.

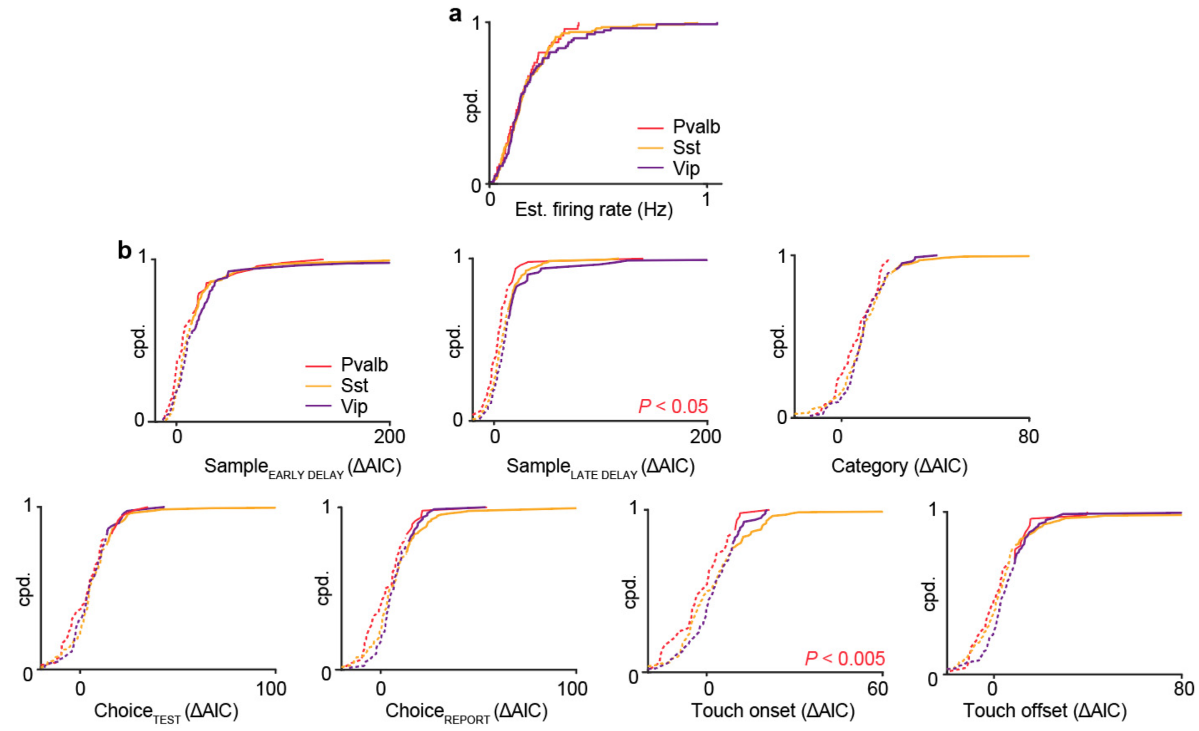


**Supplementary Figure 14. Additional task encoding across inhibitory classes. a,** Cumulative probability distributions of estimated firing rate across three inhibitory classes. **b,** Encoding strength to remaining task factors not shown in **Fig. 4** across inhibitory classes (Mann Whitney *U* test). Solid and dotted lines correspond to significant ($P < 0.01$) and non-significant encoding strengths determined via $\chi^2$ test. $n = 272$ neurons from 7 animals.

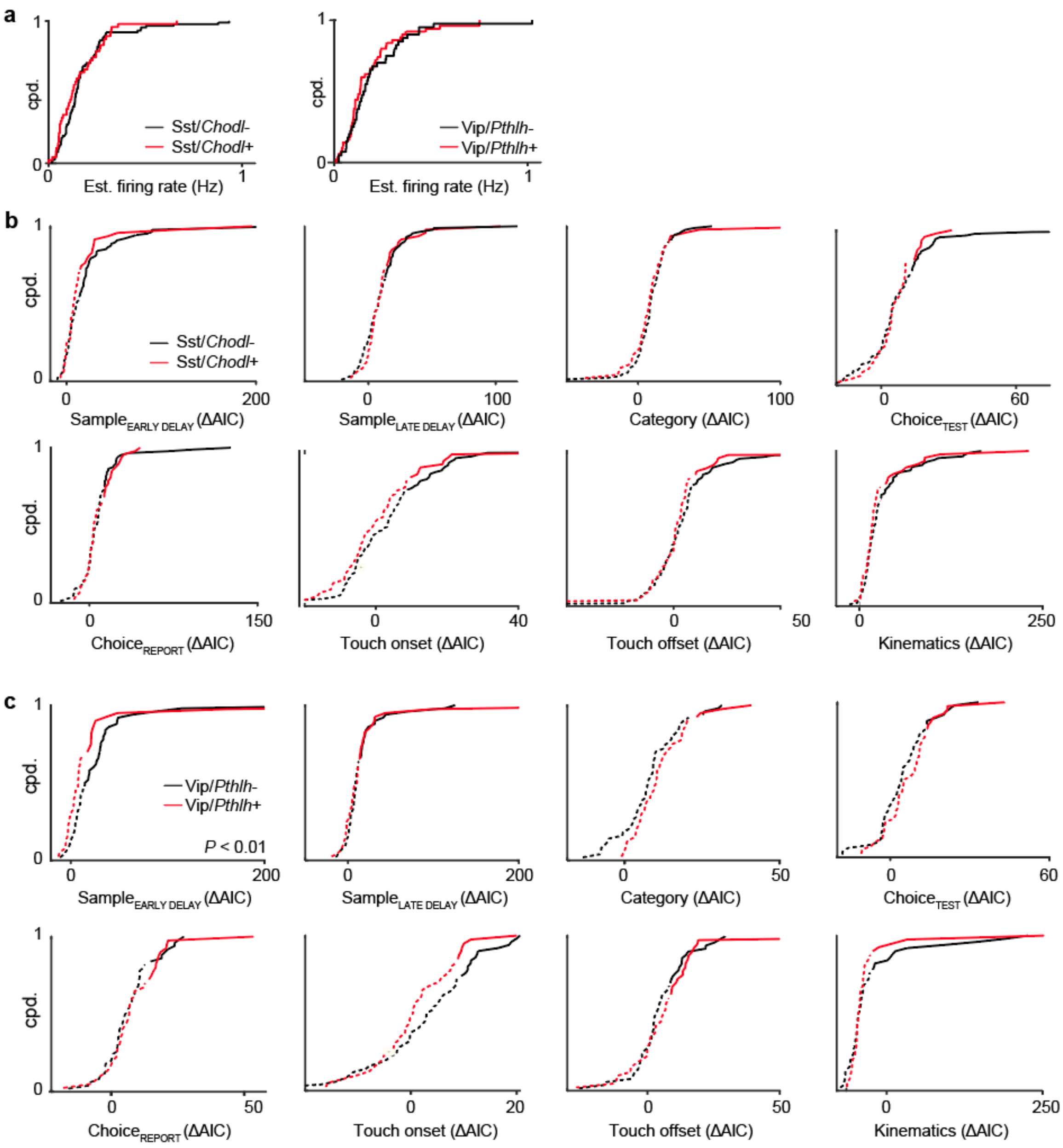


**Supplementary Figure 15. Additional task encoding across inhibitory subclasses. a,** Cumulative probability distributions of estimated firing rate across inhibitory subclasses. **b,** Encoding strength to remaining task factors not shown in **Fig. 4** across Sst subclasses. **c,** Encoding strength to remaining task factors not shown in **Fig. 4** across Vip subclasses (Mann Whitney *U* test). In **b-c,** solid and dotted lines correspond to significant ($P < 0.01$) and non-significant encoding strengths, determined via $\chi^2$ test. $n = 224$ neurons from 7 animals.

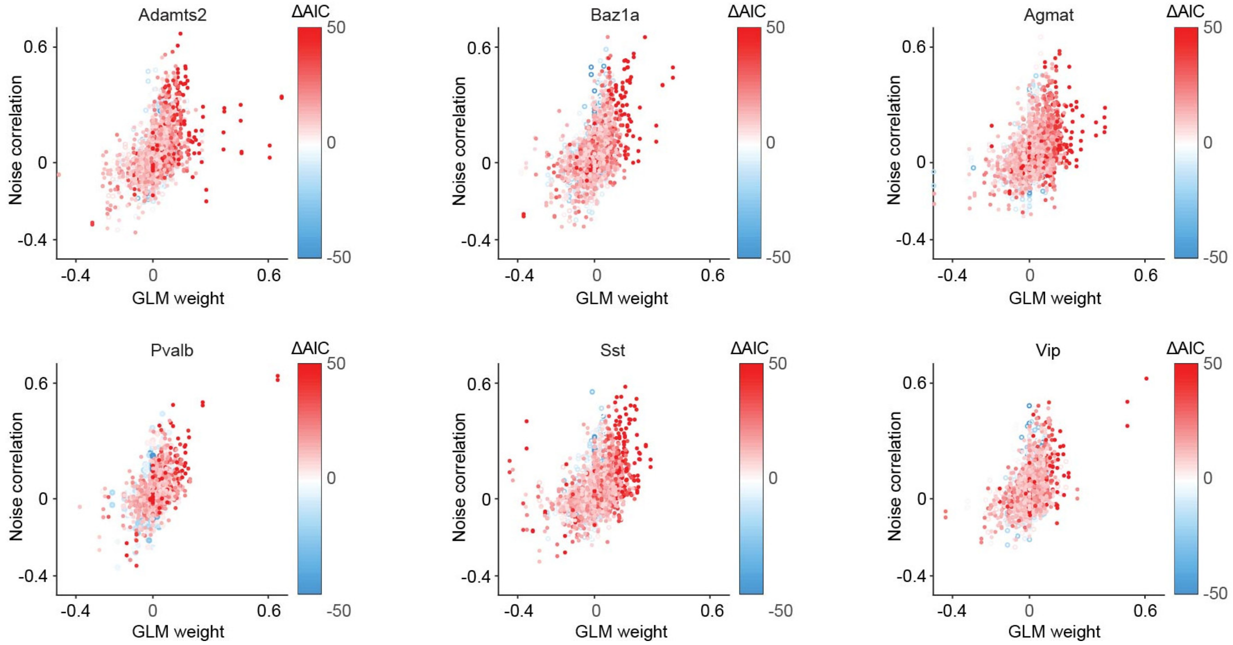


**Supplementary Figure 16. Analysis of cell type coupling factors.** For each cell type coupling factor, the relationship between an independently calculated noise correlation and the covariate weight to the cell type population activity from the GLM is shown for each neuron. The encoding strength for each coupling cell type factor is shown in color. Significant encoding strengths ($P < 0.01$; filled circle) and non-significant encoding strengths (open circle) are also shown, determined via $\chi^2$ test.

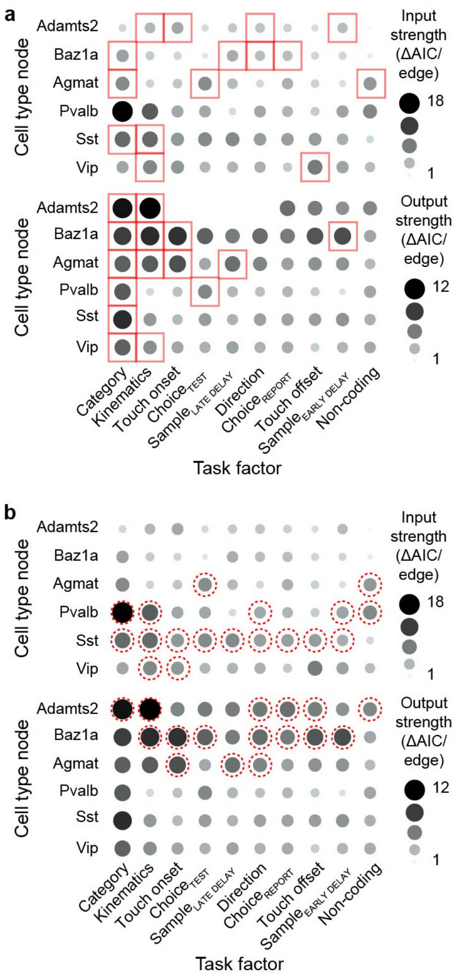


**Supplementary Figure 17. Comparison of network node strengths across and within task networks and cell types. a,** Input (top) and output (bottom) node strength for different cell types were compared across task networks. Significantly higher node strengths are noted with red box ($P < 0.05$, bootstrap test). **b,** Input and output node strength for different cell types were compared within each task network. Significantly higher node strengths are noted with red dotted circle ($P < 0.05$, bootstrap test).

### SUPPLEMENTARY NOTE

#### S.1. Sampling of Neurons with the CRACK platform

In principle, the CRACK platform offers an unbiased survey of cell type-specific functional responses across a neuronal population. In this study, task responses of neuronal cell types were restricted to neurons expressing RCaMP1.07 that were observed to be active during the imaging session. The relative number of active RCaMP1.07+ neurons in each subclass or cell type was observed to be disproportionate to previously measured cell type distributions in L2/3 S1[1-4] (**Supplementary Table 2**). RCaMP1.07 was expressed virally using an AAV with PhP.eB serotype[5]. The tropism of AAV serotypes has been shown to preferentially infect certain neuronal cell types[6]. We assessed the distribution of RCaMP1.07+ neurons in control tissue across cell types stained with HCR-FISH (**Supplementary Fig. 2**). Overall, we observed that AAV.PhP.eB had a higher likelihood of infecting excitatory neurons over inhibitory neurons. For inhibitory neurons, the tropism of the AAV.PhP.eB produced an under-representation of Pvalb and Vip class neurons compared to Sst class neurons. Whether this under-representation is stochastic or specific to certain Pvalb or Vip subtypes remains to be determined. In the future, the use of pan-neuronal transgenic lines will alleviate potential biases could result from viral tropism.

Neurons in the calcium imaging data were automatically segmented using a constrained nonnegative factorization (CNMF) approach[7]. This approach identifies neurons that exhibit active calcium responses. The activity levels of these automatically segmented neurons are shown (**Fig. 2g, Supplementary Fig. 14a, 15a**). Neurons identified as potentially inactive can be either interpreted as non-spiking neurons or those whose calcium signal-to-noise levels were not sufficient to be segmented using CNMF. The number of identified active and inactive neurons within the imaging field of view were quantified (**Supplementary Table 2**). Some cell type differences were observed. For example, Baz1a neurons were more likely to be identified as active than other excitatory cell types which is also consistent with the higher firing rates of active Ba1z1a neurons. Taken together, the results indicate choice of viral tropism and activity-based segmentation algorithms can influence sampling of neurons .

#### S.2. Comparison of Calcium Response Properties and Spike Inference across Cell Types

In the CRACK platform, two-photon calcium imaging is critical for monitoring functional responses of neurons in a manner that is compatible with post-hoc multiplexed FISH. Since calcium transients are a surrogate for spiking activity, calcium transients must be deconvolved before use in GLMs to more accurately characterize response patterns at timescales relevant to behavior. Methods to infer spikes from calcium transients using deconvolution rely on calibration experiments involving simultaneous electrophysiology and calcium imaging as well as an understanding of the response properties of the calcium indicator and biophysical properties of calcium within neurons. In this study, calcium responses were deconvolved using an Online Active Set method to Infer Spikes (OASIS), a generalization of the pool adjacent violators algorithm (PAVA) for isotonic regression[8] (**Supplementary Fig. 7a**). The extent to which the relationship between action potential and calcium events varies across cell types and how OASIS accommodates this variability is an open question. Calibration experiments in inhibitory classes using the indicator GCamp6f show that calcium responses in Pvalb neurons can underestimate spiking activity compared to Sst and Vip neurons[9,10]. Such calibration experiments are facilitated by transgenic labeling so that cell types can be selectively targeted. Currently, there are no transgenic lines that selectively label Agmat, Baz1a, and Adamts2 excitatory cell types.

Without the ability to obtain calibration data with RCamp1.07 for all cell types investigated in the study, we focused solely on assessing cell type differences in the fluorescence signals (baseline fluorescence, noise levels, rise time, and decay time) that could potentially contribute to differences in spike inference using OASIS. Baseline fluorescence intensity (bottom $10^{th}$ percentile of signal intensity) would reflect expression levels RCamp1.07 or resting calcium concentration (**Supplementary Fig. 7c**). We found no differences between excitatory cell types. For inhibitory cell classes, Pvalb showed higher relative baseline fluorescence compared to other inhibitory classes ($P < 0.05$, two-tailed Student's *t*-test with Bonferroni-Holm post-hoc correction) potentially reflecting unique buffering capacities[11]. No differences were observed in shot noise levels between cell types (**Supplementary Fig. 7d**).

In OASIS, spike estimates rely on a convolution kernel consisting of exponential rise and decay time constants that are self-tuned for each cell using an autoregressive (AR) model. Since rise and decay times are modeled from the raw calcium data for each cell, this provides a generative method based on the AR model for comparing expected calcium kinetics across cell types. No differences in rise time were observed between cell types (**Supplementary Fig. 7e**). Adamts2 cells show faster decay times than other excitatory neurons ($P < 1\text{x}10^{-5}$, two-tailed Student's *t*-test with Bonferroni-Holm post-hoc correction) while Pvalb cells exhibited faster decay times compared to other inhibitory neurons ($P < 0.05$, two-tailed Student's *t*-test with Bonferroni-Holm post-hoc correction; **Supplementary Fig. 7f**).

To determine whether cell type differences in calcium response properties influenced spike inference, we examined the distribution of spike estimates ($\hat{s}$) resulting from deconvolution. The magnitude of $\hat{s}$ relates to the confidence that $\hat{s}$ is a spiking event, with lower $\hat{s}$ values reflecting poorer estimates. While $\hat{s}$ is thresholded into binary events for downstream analysis of neuronal activity, the distribution of $\hat{s}$ extracted from a time series can be used to determine the performance of the auto-regressive model for a given cell. For each cell, we compared model performance by performing a receiver operating characteristic analysis across $\hat{s}$ values. In general, we found that lower noise levels and longer decay times were weakly correlated with better model performance (**Supplementary Fig. 7b**). However, these relationships did not necessarily translate when comparing cell type differences. For excitatory neurons, model performance was better for Agmat neurons ($P < 1\text{x}10^{-5}$, two-tailed Student's *t*-test with Bonferroni-Holm post-hoc correction) and weaker for Adamts2 neurons ($P < 1\text{x}10^{-5}$, two-tailed Student's *t*-test with Bonferroni-Holm post-hoc correction; **Supplementary Fig. 7g**). No differences were observed for inhibitory neurons. With the known exception of Pvalb neurons, we conclude that calcium response properties between cell types do not systemically differ in a manner that would result in differences amongst the other cell types when inferring spikes using OASIS.

We finally asked whether calcium response properties and spike estimates had any relationship to GLM performance. Across the analyzed neuronal population, we observed no correlation between overall GLM fit for the task encoding model and calcium signal noise level and only weak correlations when compared to calcium decay time, calcium rise time, and the spike estimate performance (**Supplementary Fig. 9a**). This demonstrates that differences in calcium responses properties from which deconvolution was performed has minimal influence on GLM performance across individual neurons that would explain task-related differences across cell types investigated.

##### S.3. Task Encoding Across Neurons

A generalized linear model (GLM) was used to assess the encoding of task-related responses in single neurons assuming a Poisson distribution of estimated deconvolved spike events. Variables representative of trial type were represented as boxcars that spanned epochs of the trial period. Whisker kinematics and population activity were represented as continuous variables. While the use of kernels to form basis functions can capture the time-varying relationship between behavior events and the neuron's probability of spiking[12,13], kernels were excluded in the GLM for a few reasons. Representing task variables through multiple kernels, substantially increases the number of covariates included in the GLM which increases the chance of model overfitting. While information about temporal response properties are not captured in the task encoding model, these dynamics are not necessary for comparing the relative strength of task-related responses between cell types. Rather, the exclusion of kernel reduces model complexity and eases the interpretation of the model results.

The information content of a neuron's activity for a given task factor was determined by comparing the AIC of the full model containing the task factor against the AIC of the partial model which excluded covariates representing that task factor. We chose to use AIC to convey GLM fit because it is a principled way to compare fit between models of different complexity (i.e. different number of model parameters). AIC accounts for the trade-off between the goodness of fit (risk of overfitting with large number of model parameters) and simplicity (risk of underfitting with too few model parameters). ΔAIC reports the difference in model fit between a full model and partial model that is somewhat agnostic to model complexity. Statistical significance and ranking of models can be assessed using ΔAIC values, which enables information content to be compared both within and across different task factors and cell types.

It should be noted that ΔAIC differs from deviance explained, a measure of the goodness of model fit by comparing the full model to the null model. To illustrate this, we compared deviance explained for the full model against the ΔAIC of each task factor for each neuron (**Supplementary Fig. 9b**). The majority of task factors were positively correlated with model fit. In particular, strong correlations with stimulus direction, whisker kinematics, and population coupling suggest that these task factors were key drivers of goodness of fit for the GLM used in this study. However, task factors for category and touch offset showed no significant correlation. This demonstrates that information content of a given task factor and goodness of model fit are not necessarily related. A neuron's activity can encode information about a given task factor while simultaneously having a poor overall model fit.

##### S.4. Measuring Functional Connectivity using Cell Type Coupling

In order to assess how a neuron's activity relates to the surrounding population, the dimensionality of the population activity was reduced using non-negative matrix factorization as previously described[13]. For analysis of population coupling in the task encoding GLM, the population activity was factorized into varying ranks. Ranks were defined as an individual covariate in the model and characterized as the functional response of non-overlapping subpopulations. The task factor for coupling consists of all of these rank covariates. We measured coupling as increasing number of ranks are included in the model (**Fig. 5a**) and observed that ΔAIC increases with the number of ranks. This means that as more of the variance in the population activity is captured in the GLM, neuron's activity can be better explained by the activity from these different subpopulations. For the cell type coupling GLM, the population activity was subdivided by cell type. Since subdividing by cell type reduces the number of neurons in each subpopulation, a factorization rank (r = 1) was used to standardize the population activity of each cell type.

Co-fluctuations in activity between neurons while controlling for stimulus and other task conditions ('noise correlations') are often used as a measure of functional connectivity of neurons[14]. In the cell type coupling GLM, we fit a neuron's activity to task variables as well as the cell type coupling covariates capturing the activity to each of the simultaneously recorded cell types. The covariate weights obtained in the GLM are similar to noise correlations. For illustration, we plot the weights of cell type coupling covariates for neurons that encode stimulus direction against the noise correlation across trials with the same stimulus condition (**Supplementary Fig. 9**). As can be seen, positive or negative covariate weights typically correspond to positive or negative noise correlations.

Functional connectivity of a cell type is computed as the ΔAIC of the full model against the partial model excluding the cell type coupling factor. This calculation asks whether the co-fluctuations in activity that are observed can be explained by activity related to task variables or that of other neurons recorded in the population. Thus, a significant, positive ΔAIC identifies noise correlations that are specific to that cell type and cannot be explained by other recorded neurons. Positive or negative correlations can both produce a significant, positive ΔAIC.

A significant, positive ΔAIC value of a neuron's cell type coupling factor serves to reject the hypothesis that those input sources are not shared by the other recorded neurons since they would have been explained away by the other covariates. It can also raise two possible interpretations. The first interpretation is the presence of a direct synaptic interaction between the neuron and the cell type. The second interpretation is that the neuron and the cell type share common input from a source that is not captured in the population recording. This is because the ΔAIC can only explain the activity patterns from simultaneously recorded neurons included in the full model. This second interpretation may be applicable to functional connectivity between inhibitory cell types in which significant coupling weights are not negative as would be predicted from direct inhibition. Positive weights when considering inhibitory-inhibitory and excitatory-inhibitory functional connectivity could potentially be explained by excitatory drive from a non-measured source such as bottom-up input from L4 or thalamus or top-down input from high-order areas.

Functional connectivity analysis provides complementary insight into how cell types interact under task conditions. Our analysis of task networks show that cell types interactions are layered upon coding features of individual cells. Certain task-related features (ex. category and whisker kinematics) involve strong local network interactions while other features ($\text{sample}_{\text{LATE DELAY}}$) may be inherited from other areas[15]. Labeling connections as strong and stable vs. weak and variable provides clues as to which connections may be stable, intrinsic motifs vs. transient, dynamic interactions. Ultimately, distinguishing between direct and indirect connectivity requires confirmation experiments to test synaptic connectivity (**Fig. 6**).

#### SUPPLEMENTARY NOTE REFERENCES
